# A gene-distal polygenic architecture underlies the dominant axis of human trait covariation

**DOI:** 10.64898/2026.05.26.728050

**Authors:** Olin K. Silander

## Abstract

Genome-wide significant loci explain only part of the heritable signal for most complex traits, leaving a broad sub-significant polygenic background whose organisation is largely uncharacterised. Whether this background is largely diffuse noise or contains coherent structure shared across traits has been difficult to test, because SNP-level effects are noisy and LD-correlated, and conventional cross-trait analyses have exclusively focused on GWAS significance-filtered, LD-pruned variants. Here, using 403 GWAS summary statistics, we show that coherent cross-trait signal is detectable using signed regional medians of SNP effects. Singular value decomposition of regionally averaged effects recovered latent phenotypic axes. Focusing on the sub-significant polygenic background, we found that the axes persisted after removal of all genome-wide significant loci and flanking regions observed in any of the 403 GWAS analysed, a step which eliminated more than 75% of genomic loci. The dominant axis of trait variation, primarily metabolic, but spanning anthropometric, musculoskeletal, and cognitive traits, was unlike secondary axes in being enriched for regions more than 250 kb from any gene body, both before and after removal of genome-wide significant loci observed in these GWAS. These results show that the sub-significant polygenic background carries structured cross-trait information, and identify gene-distal non-coding variation as a defining feature of the dominant axis of human trait covariation, patterns that are largely inaccessible to approaches that rely on genome-wide significance for variant prioritisation.

## Introduction

Genome-wide significant loci explain only part of the heritable signal for most complex human traits, leaving a broad sub-significant polygenic background whose organisation remains poorly understood. This background is difficult to analyse directly: SNP-level effects are noisy, strongly correlated through LD, and individually weak (Zhang et al. 2018; Visscher et al. 2021; Fisher 1919; Boyle et al. 2017; Reich et al. 2001). At the same time, complex traits are not genetically independent, complicating interpretations. Many traits share regulatory programmes, cell types, developmental processes, and molecular pathways (Calderon et al. 2017; Kim et al. 2025; GTEx Consortium 2020; Finucane et al. 2018), and individual loci often have pleiotropic or correlated effects across phenotypes (Jee et al. 2026; Watanabe et al. 2019; Zhang et al. 2023). Together, these two factors, pervasive pleiotropy and pervasive polygenicity, have limited our understanding of the genetic architecture underlying human trait variation.

Latent decomposition methods provide a direct way to study shared genetic structure across traits. By decomposing matrices of GWAS summary statistics across large phenotype panels, these approaches have recovered axes of phenotypic organisation and pleiotropic genetic effects (Tanigawa et al. 2019; Zhang et al. 2023; Lazarev et al. 2025; Backman et al. 2021; Jee et al. 2026; Oblong et al. 2024). However, the preprocessing steps used by many of these methods are not neutral for the question of whether sub-significant GWAS loci impart trait structure. Significance-based filtering, LD pruning, or index SNP selection before decomposition (Trevisan et al. 2025; Tanigawa et al. 2019; Oblong et al. 2024) reduce the genetic variation space, reducing sensitivity toward polygenic effects, and biasing analyses toward large effect variants, which are often enriched near genes (Spence et al. 2026; Watanabe et al. 2019), and concomitantly often depleted in intergenic regions relative to null expectations. As a result, these methods can recover coherent latent axes, but cannot directly test whether similar axes persist in sub-significant regions, or how that architecture is organised.

However, other approaches do incorporate sub-significant GWAS signals. LDSC and S-LDSC (Bulik-Sullivan et al. 2015; Finucane et al. 2015) use genome-wide information, and cross-trait LDSC estimates genetic correlations across traits. However, these approaches summarise pleiotropy as genome-wide averages, or as averages within predefined functional annotations, with no genomic localisation. Genomic SEM (Grotzinger et al. 2019, 2026) explicitly models latent genetic trait covariance, but typically uses a pre-specified phenotypic covariance structure. Locus-based approaches such as LAVA (Werme et al. 2022) retain sub-significant signal and genomic localisation, but are designed to test local heritability and genetic correlation within predefined loci, rather than to discover genome-wide latent axes across many traits.

Here we take a different approach. We aggregate SNP-level effects into regional summaries to obtain a matrix of regional genetic effects across multiple GWAS traits. We then apply SVD to this region-by-trait matrix, without genome-wide significance filtering, LD pruning, or index SNP selection. Regional median aggregation reduces SNP-level noise, is robust to outlier effects, and compresses local LD-correlated effects into regional summaries, while retaining the signed direction of genetic effects. This aggregation does not assume that all causal variants in a region act in the same direction. Rather, it captures regional signed imbalance, and is therefore most sensitive to regions where the distribution of signed SNP effects is shifted away from zero.

We show that SVD of the region-by-trait matrix recovers coherent latent axes of phenotypic organisation, and that retaining the signed directionality of effects is necessary: randomising the signs of window median effects fully ablates the recovered axes, and ignoring signs substantially reduces latent axis coherence. The latent axes persist after removal of all genome-wide significant loci across all GWAS we considered, directly demonstrating that sub-significant polygenic signals carry coordinated, signed, cross-trait structure. Most notably, the dominant latent axis, primarily metabolic but spanning anthropometric, musculoskeletal, and cognitive traits, exhibits a gene-distal polygenic architecture that persists regardless of whether genome-wide significant loci are included. This distinguishes the primary axis from secondary axes and implicates gene-distal regulatory variation as a major feature of the broadest dimension of human phenotypic covariation.

## Results

### Regional effect aggregation reveals latent phenotypic axes

We selected 403 high-quality GWAS studies spanning 16 broad phenotype categories (Suppl. table 1) from the OpenGWAS database. For each GWAS, we summarised genome-wide genetic effects as the median signed beta across SNPs within non-overlapping 50 kb genomic windows (minimum 30 SNPs per window (Suppl. Fig. 1)). This regional aggregation reduces dimensionality and mitigates noise inherent in less common or small-effect variants, without selecting variants based on significance or reducing each locus to an index SNP and effect. SVD of the resulting 403 x 51,907 matrix revealed clear low-dimensional trait structure, with robust leading components (Suppl. fig. 2a-d) capturing coherent groupings of biologically related traits (Fig. 1a–c). PC1 loaded broadly across metabolic, anthropometric, musculoskeletal, and cognitive traits, whereas later PCs were more closely associated with individual trait categories.

**Figure 1.**
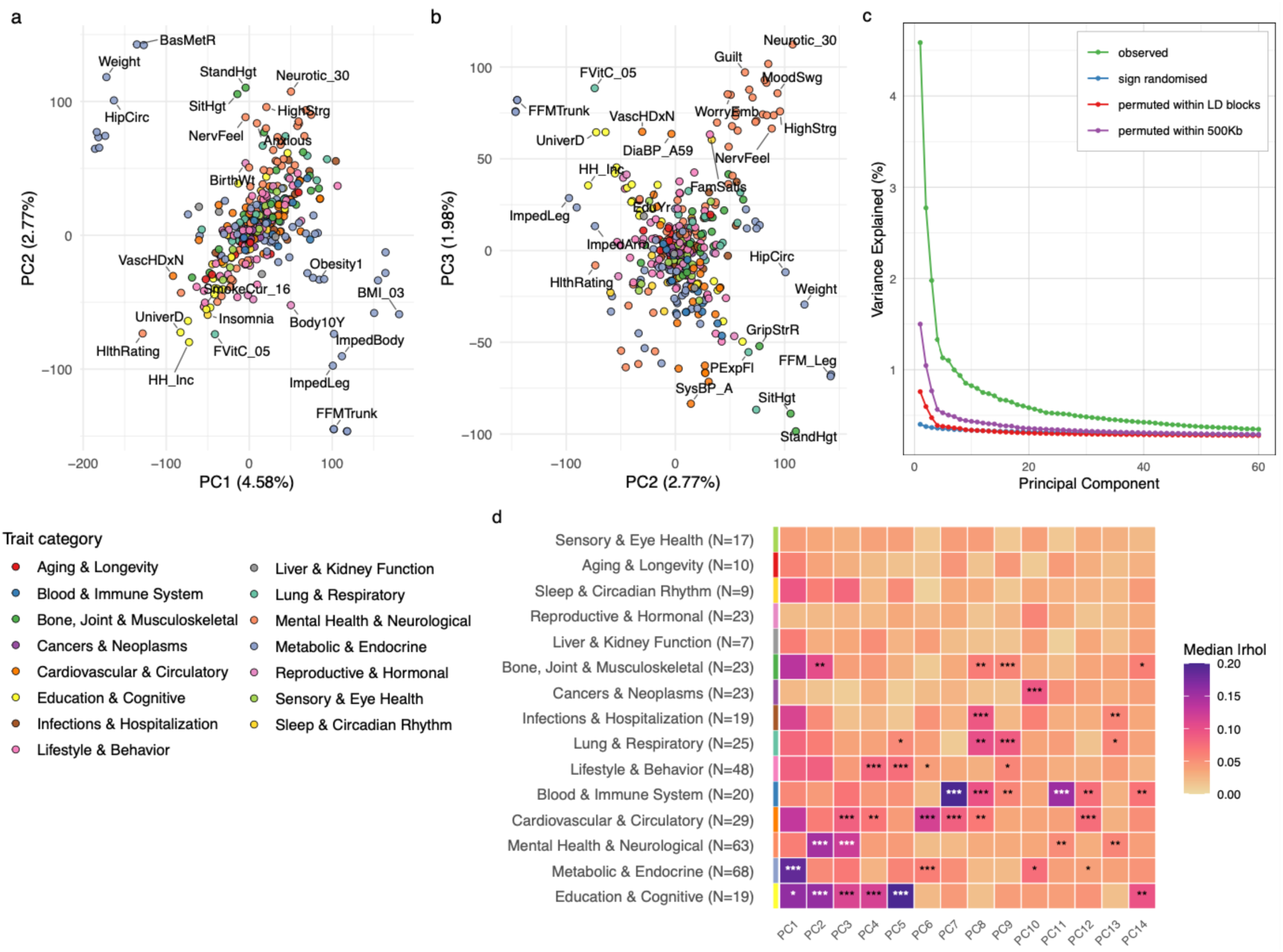
Regional aggregation of signed GWAS effects recovers coherent latent phenotypic axes. **a**. Trait projections on PCs 1 and 2. Points are coloured by phenotypic category. **b**. Trait projections on PCs 2 and 3. **c**. Scree plot of explained variance for the observed data and permuted datasets. **d**. Median absolute Spearman correlations of traits within each phenotype group across PCs. Colours correspond to the strength of the median correlation within a phenotype category. Asterisks indicate the significance of correlations in that group, as determined by 1,000 label permutations across groups, and ranking. Statistical significance within a category is affected by variation in the strength of correlations for phenotypes in that category (i.e. high within-category variance in correlations). *** p < 0.001, ** p < 0.01, * p < 0.05.

For example, PC2 and PC3 were highly correlated with mental health and neurological traits; PC6 with metabolic and cardiovascular traits; PC10 with cancers and neoplasms; and PC11 with blood and immune traits (Fig. 1d, Suppl. figs. 2-6). The SVD yielded window-level loadings identifying genomic regions driving each latent axis. We observed that strongly loaded windows on different PCs were enriched for specific functions. For example, PCs 1-5 and 10 were enriched for genic regions with functions relating to neuronal function, and PC9, an immune-respiratory axis, was enriched for functions relating to T cell differentiation (Suppl. fig. 7).

This latent structure depended critically on the directionality and local structure of effects. Replacing SNP betas with their absolute values, randomising signs across the matrix, increasing the genomic window size beyond 500 kb, or permuting window locations within LD blocks each fully ablated or substantially diminished the signal (Fig. 1c; Suppl. figs. 8-12). This indicated that the recovered latent structure was not an artefact solely of effect size distributions or genomic block structure. Rather, we hypothesised that regional median aggregation denoises effect estimates such that the window median reliably reflects weak causal signal due to single or multiple variants, partially mediated by LD, even when no individual SNP reaches significance. SVD identifies windows where this directional signal is consistent across traits, recovering cross-trait structure invisible to single-locus analysis.

Genomic loadings driving latent phenotypic axes are localised and PC-specific In many, but not all cases, the loadings of the loci on the PCs tended to consist of single isolated peaks (Fig. 2a-c; Suppl. figs. 13-15). Overall, they exhibited significantly more peaked and extreme loadings than expected (permutation test, p < 0.01; Suppl. fig. 14). Several high-loading windows corresponded to established pleiotropic loci (Suppl. fig. 15). For example, a single window within the Chr16 FTO locus (with known pleiotropic effects across metabolic and neurological domains) was heavily loaded for PCs 1 and 2 (metabolic and cognitive/mental health axes; Figs. 1, 2a, and 2d); several neighbouring regions were heavily loaded at the Chr11 NCAM1 locus for PCs 2-4 (Fig. 2e); and for PCs 2 and 3, several adjacent windows loaded heavily at the Chr4 LCORL locus (Fig. 2; Suppl. fig. 15). In general, strongly loaded windows were proximal to known GWAS loci (chromosome-matched permutation test, p < 0.001; Suppl. fig. 16). However, other strongly loaded regions had no significant GWAS signal in any of the 403 studies here, nor did they have obvious correspondence to other known GWAS loci, including PC3 and PC4 loadings within the CNTLN/SH3GL2 locus at Chr9 17.2 Mb; or PC2 and PC3 at the C17orf51 locus at Chr17 21.45 Mb (Fig. 2c and 2f; Suppl. fig. 17), showing that the axes are not simply a recapitulation of known associations. This suggested that the latent structure was not necessarily dependent on genome-wide significant loci.

**Figure 2.**
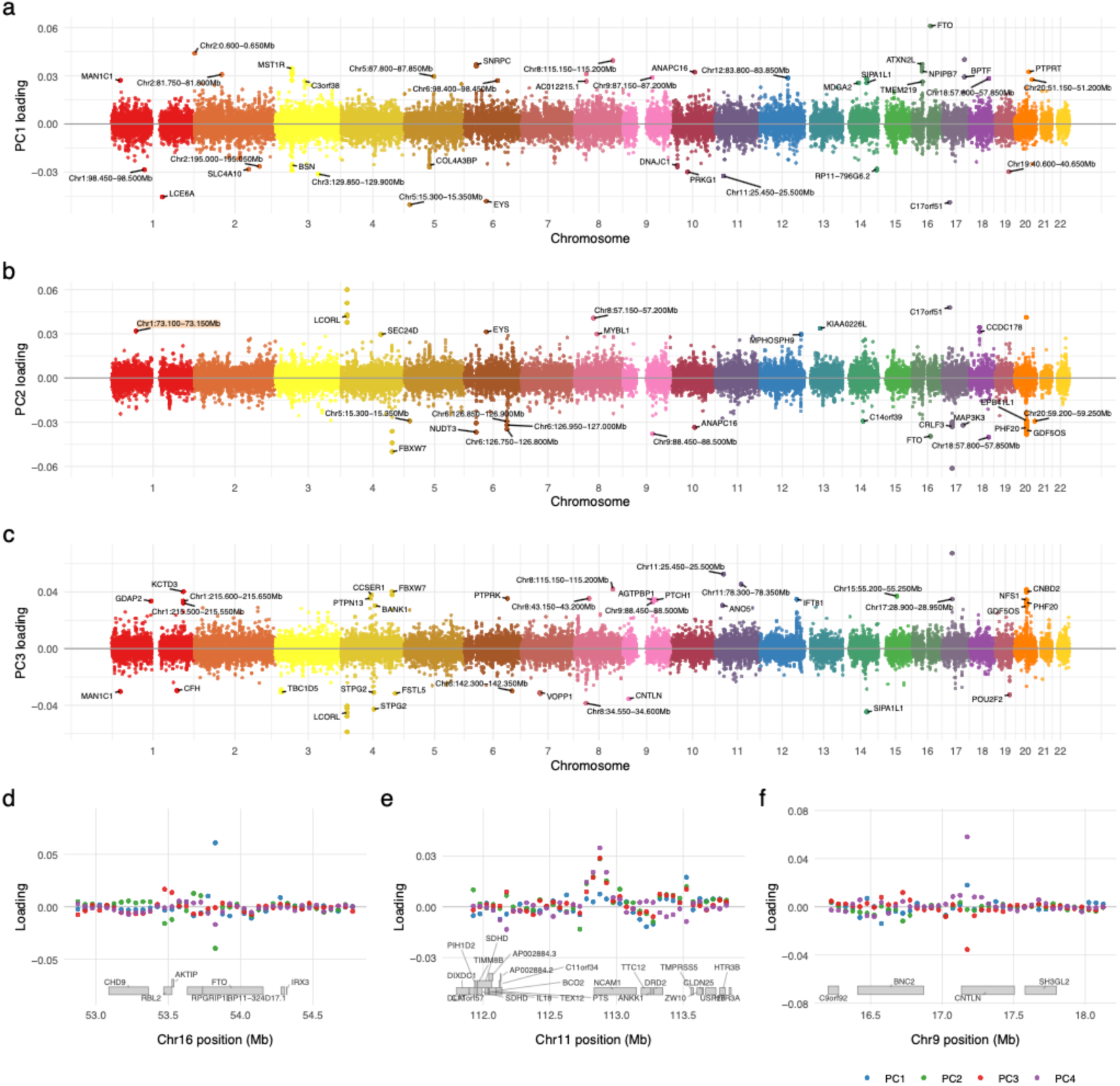
Locus loadings exist as sparse peaks distributed across the genome. **a-c.** Miami plots of PC loadings per window, arranged by chromosome. The top 0.1% strongest-loaded loci are labelled; when multiple neighbouring windows are among the top peaks, only one is labelled. **d-f**. Regional zooms showing PC loadings across the first four PCs at each region. The loading on each of the first four PCs is indicated. **d.** FTO at the chromosome 16 53-54.5 Mb locus. **e.** NCAM1 at the chromosome 11 112-113.5 Mb locus. **f.** CNTLN/SH3GL2 at the chromosome 9 16.5-18 Mb locus.

### Sub-significant polygenic effects drive coherent latent phenotypic axes

To directly test this, we removed all genomic windows containing any SNP significant at p < 5e-8 across any of the 403 studies. To mitigate the effects of LD, we also removed 100 kb of flanking sequence on each side (two 50 kb windows), reducing the matrix from 51,907 to 12,688 windows, a reduction of more than 75% (Suppl. fig. 18a), and removing up to 90% of the most strongly loaded loci (Suppl. fig. 18b). Despite this extensive locus pruning, the latent axes were reproduced with high fidelity (Figs. 3a–c): Procrustes similarity between the full matrix and GWAS-depleted latent trait configurations was 0.92 using the first four PCs and 0.88 for the first eight (Suppl. fig. 12); trait-category correlations with PCs were well preserved (Suppl. fig. 18c); and window loadings on the retained non-GWAS windows were strongly correlated between the full and GWAS-depleted SVD (PC1 r=0.98; Suppl. Fig. 19), declining across later components, as expected. This demonstrated that the latent structure is not fully dependent on established GWAS loci.

**Figure 3.**
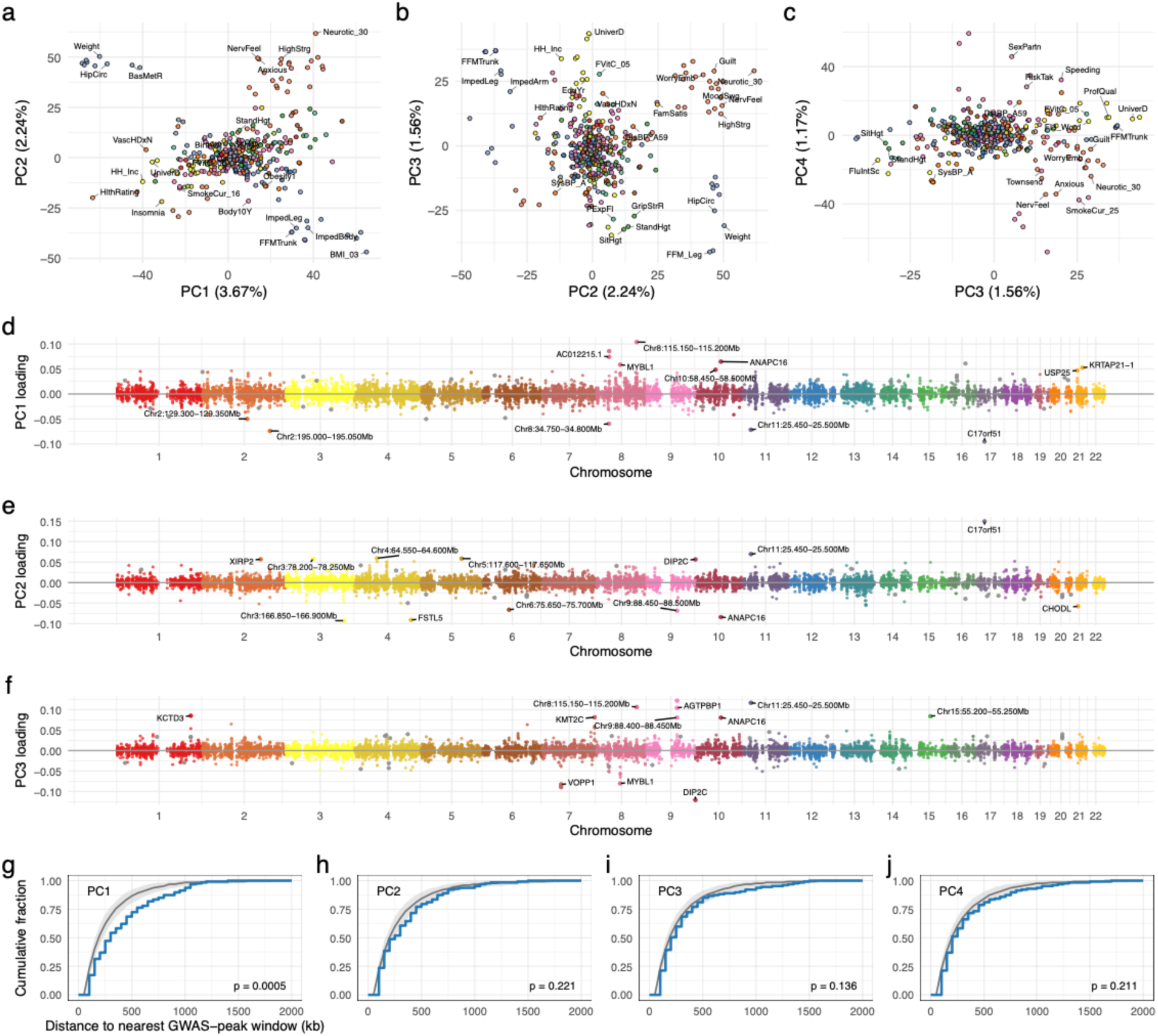
Latent axes persist after removal of GWAS-significant loci. **a-c.** Trait projections on PC pairs. **d-f.** Miami plots after removal of GWAS-significant loci. The top 0.1% most strongly loaded loci are labelled. For adjacent strongly-loaded loci, only the central window is labelled. **g-j.** Cumulative frequency plots showing the distance from the top 1% most strongly loaded windows to the nearest GWAS-removed window. The grey ribbon shows the median and 95% null interval based on chromosome-matched random window sets; the observed distribution is shown in blue. The p-values shown on the bottom right of each panel are from a permutation test comparing the area between the observed CDF and the mean null CDF using 2,000 permutations.

In addition, the functional enrichment patterns we observed in the full dataset were similar to those in the GWAS-depleted dataset, with PCs 2 and 4 strongly enriched for neuronal functions, and secondarily PCs 3, 7, and 11. However, and in strong contrast to the previous results, we observed no strong functional enrichment of any category on PC1 (Suppl. Fig. 20).

Similarly, we observed many isolated windows with strong loadings (Fig. 3d-f), as before exhibiting more extreme and more localised values than expected (p < 0.01, permutation test; Suppl. fig. 21), although again not all regions exhibited this pattern (Suppl. fig. 22). Importantly, we found that the rank-importance of retained individual windows shifted substantially after GWAS removal (Suppl. Fig. 23), revealing new high-loading regions, consistent with the sub-significant polygenic signal being distributed across regulatory regions outside the current GWAS catalogue.

As a further control against this result arising from LD leakage from removed loci, we tested whether top-loaded windows in the GWAS-depleted SVD were more proximal than expected to the GWAS-removed regions. For PC1, top-loaded windows were in fact significantly more distal from GWAS-removed regions than chromosome-matched nulls (Fig. 3g; p < 0.0005, permutation test), while for PCs 2-4 they were no closer than expected from null permutations (Figs. 3h-j). This suggested LD leakage was not a significant explanatory phenomenon for the recovered latent structure.

### The primary latent axis is characterised by gene-distal variation

Having established that retained sub-significant windows carry strong latent structure, we sought to understand the genetic architecture underlying the latent axes. We thus examined the genomic localisation of the loci driving each axis. We calculated the proximity of strongly loaded loci to genic and non-genic regions for PCs 1-4. When considering all loci (i.e. including GWAS-significant loci), PCs 2-4 showed strong enrichment for genic regions among top-loading windows (1.11 - 1.20-fold enrichment of windows directly overlapping gene bodies; p ≤ 0.005, permutation test; Fig. 4a and Suppl. fig. 24), consistent with the known gene-proximity of GWAS-significant variants. PC1 showed a distinctive pattern: top-loaded windows were modestly enriched in genic regions (maximum 1.09-fold; p = 0.025, permutation test; Fig. 4a; Suppl. fig. 24), and enriched at large distances from any gene body, apparent as a significant shift in the cumulative distance ranked curve (p = 0.0104, permutation test; Fig. 4b) and a strong fold-change (>250 kb; 1.27-fold, p = 0.023 for the top 1%; 1.31-fold, p = 0.001 for top 2%; Figs. 4a, 4b), relative to chromosome-matched nulls. There was no such distal region enrichment for PCs 2-4 (Suppl. fig. 25), which were in fact distal-depleted in almost all cases (Suppl. fig. 24). Concomitantly, from a functional perspective, PC1 was not enriched in promoter regions, and not depleted in intergenic regions, in contrast to PCs 2-4 (Fig. 4c). This pattern indicated that the dominant latent axis uniquely exhibited a distinct gene-distal regulatory architecture.

**Figure 4.**
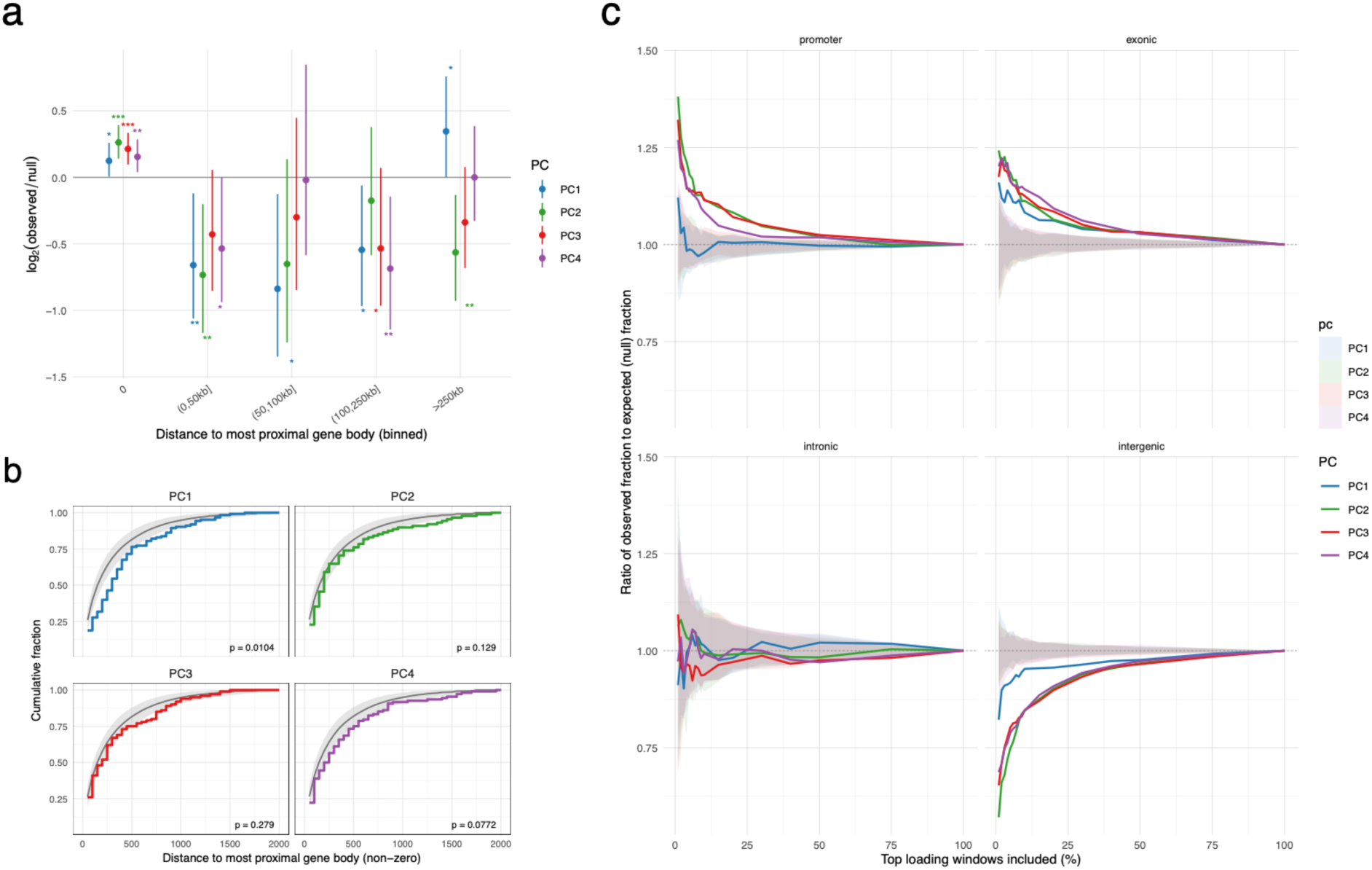
The dominant latent trait axis is enriched for gene-distal loci. **a.** Log2 fold-change in gene proximity for the top 1% most strongly loaded windows relative to chromosome-matched nulls. All PCs exhibit enrichment for genic overlaps; only PC1 exhibits enrichment for distal regions. **b.** Cumulative distances to the nearest gene-body, excluding direct genic overlaps. PC1 exhibits a clear and distinct rightward shift toward loci being more gene-distal. The p-values for a permutation test of a shift in the distribution are shown on each panel. **c.** Enrichment of functional annotation classes among top-loaded windows. For each PC, windows are ranked by absolute loading, and the ratio of observed to genome-wide overlap fraction is plotted cumulatively from the top 1% to all windows. Shaded ribbons indicate the 95% null interval across 100 chromosome-matched random sets. A ratio of 1 (dotted line) indicates no enrichment above the genome-wide background. PC1 again is distinct in enrichment and depletion patterns, showing no enrichment for promoters, little enrichment for exonic regions, and weak depletion for intergenic regions. Functional classes are mutually exclusive: Intergenic regions are defined by having no overlap with any gene body or promoter; intronic regions overlap a gene body but not an exon; exonic regions overlap an exon but not a promoter; promoters are regions ±2 kb from any transcription start site.

Repeating this analysis for the GWAS-depleted dataset, there was no consistent pattern of genic enrichment for PCs 1-4 compared to chromosome-matched nulls (top 1% of loci: fold range 0.85 - 1.20, all p > 0.05; Suppl. fig. 26). In contrast, PC1 retained a strong and consistent enrichment for loci distal to gene bodies across top-loading thresholds and flank definitions (Suppl. fig. 26 and Suppl. fig. 27). PCs 2 and 4 showed weaker and less consistent distal enrichment, detectable only under some thresholds, while PC3 showed no distal enrichment across any threshold examined. Thus, PC1 was uniquely distinguished by the strength and persistence of its gene-distal architecture across both the full and GWAS-depleted analyses. This identified gene-distal regulatory variation as a primary feature of the dominant phenotypic axis. However, distal regulatory signals may contribute to some secondary axes, most apparent when more weakly contributing loci were considered (e.g. the top 5%).

## Discussion

These results establish two fundamental architectural characteristics of human trait co-variation. First, the sub-significant polygenic background carries coherent cross-trait directional structure that mirrors, but is independent of, genome-wide significant loci. Second, and most notably, the dominant latent axis of human trait covariation has a gene-distal polygenic architecture that is present in both significant and sub-significant sequence, and is considerably stronger and more consistent than for any secondary axes. Gene-distal enrichment of this kind has been previously proposed for individual complex traits including schizophrenia and height (Loh et al. 2015; Boyle et al. 2017; Finucane et al. 2015; Maurano et al. 2012); our results suggest it is a general property of the primary axis of human phenotypic covariation, recoverable either through approaches that preserve the full polygenic signal, or from sub-significant signal alone. This sub-significant signal is difficult to detect using more common significance-based approaches. Regional averaging without significance filtering, as we have done here, mitigates the frequency bias inherent in index-SNP-based approaches: lower-frequency variants are underrepresented in p-value-based catalogues regardless of their true effect size, such that their contributions to trait architecture can be systematically lost.

The primary axis (PC1) runs across multiple trait categories and captures variation in metabolic, anthropometric, musculoskeletal, and cognitive phenotypes. This is consistent with regulatory variation in shared developmental or physiological networks that have broad downstream consequences, rather than being trait-specific. It also matches predictions of the omnigenic model: that peripheral regulatory variation propagates effects across traits through shared networks. The absence of GO enrichment for PC1 in the GWAS-depleted analysis is consistent with this interpretation, as nearest-gene annotation could systematically misassign distal regulatory elements acting over tens or hundreds of kilobases.

Several caveats apply to the results here. Most critically, the set of loci we have classified here as sub-significant was determined by the studies considered. In addition, the sub-significant nature is affected by study size, population, and error. Other genomic regions may have strong effects on other traits, and those may be correlated with the traits we consider here; sub-significant is thus only a comparative descriptor. Partial sample overlap across UK Biobank-derived studies induces correlated estimation error, though this is a global property of the decomposition and is unlikely to explain the PC1-specific spatial pattern. Regional differences in LD structure and SNP density between genic and gene-distal regions could in principle influence the stability of regional median statistics, but again would be expected to affect all PCs similarly. Our analyses are restricted to European-ancestry populations; whether the identified axes and their genomic localisation generalise to other ancestries remains to be established. More broadly, the gene-distal windows driving PC1 remain functionally uncharacterised beyond simple enrichment patterns such as neuronal processes; resolving their regulatory targets and mechanisms will require chromatin contact data and functional annotation beyond the scope of the present analysis, and suggest a natural and necessary extension of these findings.

## Methods

### GWAS summary statistics

We downloaded GWAS summary statistics from the IEU OpenGWAS database (Elsworth et al. 2020), which includes both IEU-processed studies and UK Biobank phenotypes processed through the MRC-IEU GWAS pipeline. We excluded studies with heritability score p-values greater than 1e-5; studies flagged as isBadPower, isExtremeSE, isHighSE, isLowNeff, isMidNeff, or isSexBias; protein-level and metabolic-level phenotypes (met- and prot- prefixes); traits unlikely to reflect direct genetic determination, such as cereal preference or workplace noise; and studies with fewer than 1.8M SNPs. We additionally filtered studies based on median absolute effect size, retaining only those with median |beta| between 1e-4 and 0.5, excluding studies with negligible signal or implausibly large effect sizes. This yielded 408 studies. We then examined the minor allele frequency distribution of SNPs in each study using either annotated allele frequencies present in the raw downloaded data, or allele frequencies obtained from gnomAD (Gudmundsson et al. 2022), removing five studies that appeared to have been restricted to common variants (few or no SNPs with MAF < 0.2), leaving a final set of 403 studies (Suppl. table 1).

### Exclusion of high-LD regions and sex chromosomes

To mitigate the influence of a small number of loci on the patterns of phenotypic variation, we excluded three high-LD loci with well-established large and pleiotropic effects on traits: the 17q21.31 inversion block (GRCh37 chr17:43.500 - 45.000 Mb) (Steinberg et al. 2012); the MHC locus, chr6:25.000 Mb - 34.000 Mb; and the 12q24 locus, chr12:111.200 Mb - 113.400 Mb (Scherer et al. 2006; Yu et al. 2005; Auburger et al. 2014). In addition, we excluded all sex-chromosome loci.

### GWAS effect aggregation

We partitioned the genome into fixed non-overlapping windows at regional scales: 25 kb, 50 kb, 100 kb, 200 kb, 500 kb, and 1 Mb, and calculated the median signed beta value for all SNPs within the window, inclusive of any imputed SNPs, using BEDtools (Quinlan and Hall 2010). We reasoned that regional median aggregation could reduce SNP-level noise while preserving weak regional directional structure, even within regions containing only sub-significant SNPs. SNPs within a genomic window are not independent, and groups of SNPs in LD with trait-associated variation can produce non-random signed effect patterns. When signed beta effects are coherent within a locus, the median is displaced from zero, even if no individual SNP reaches genome-wide significance, as significance is a function of effect size, allele frequency, and measurement error. Conversely, regions with opposing local effects, or with true signal confined to only a small fraction of SNPs, have medians close to zero, and this regional statistic assigns such regions as having minimal effects. The statistic is therefore a measure of regional signed imbalance, and not strongly influenced by single large-effect SNPs, as a mean would be. In addition, by aggregating across all SNPs in a window, rather than selecting the most significant index SNP, the statistic is decoupled from significance-based selection, and should be less sensitive to winner’s curse. This contrasts with index-SNP-based summaries, which can be influenced by allele frequency and local LD structure, and which is more susceptible to winner’s curse.

For each trait, we removed any windows not containing at least 30 SNPs (20 SNPs in the case of 25 kb windows). On average, each window contained 175 SNPs (Suppl. fig 1). We also removed any genomic window that was absent in more than five traits (for example due to SNP-based filtering, lack of imputation, or being centromeric). For those windows absent from five or fewer traits, the missing value in those traits was imputed as the median effect for that trait across all retained windows. These exclusions resulted in a final matrix of 51,907 genomic windows and 403 traits.

### Trait Categorization

Traits were assigned to broad phenotype categories using LLM-assisted classification. Trait names and descriptions from the IEU GWAS database were provided to the model with instructions to propose biologically coherent groupings, evaluate internal consistency, and revise category boundaries iteratively. The resulting assignments were reviewed manually; one trait (professional qualifications: nursing, teaching) was reassigned from Lifestyle and Behaviour to Education and Cognitive. The final categorical assignments are provided in Suppl. table 1.

### Singular value decomposition

We applied SVD to the 403-trait by 51,907-window effect matrix, standardised by centering and scaling each trait’s effect vector to mean zero and unit variance across windows. Without scaling, the decomposition would be dominated by traits with the largest marginal effect sizes, which set the leading components regardless of whether their directional pattern is shared across traits. Scaling to unit variance allowed each trait equal geometric weight, with the SVD finding axes along which many traits are coherently aligned, rather than axes on which traits have large effects. We used the prcomp_irlba function from the irlba R package v2.3.7 (Lewis 2017).

This decomposition yields window-level loading vectors and trait-level score vectors, which we refer to as PC loadings and trait scores respectively. Importantly, trait encoding directions in IEU GWAS, which are arbitrary, should not affect window-level PC loadings as flipping the sign of any trait’s effect vector does not affect the window loading vectors, only the corresponding entries in the trait score matrix. Thus, whether a trait is encoded as fat mass or fat-free mass, the window loading structure is identical; only the sign of that trait’s projection onto each PC differs.

### PC orientation

Because the sign of each PC is arbitrary, we oriented each component by a pre-specified anchor trait: BMI_03 (BMI) for PC1, standing height (StandHgt) for PC2, neuroticism score (Neurotic_30) for PC3, and diastolic blood pressure (DiaBP_A59) for PC4, each of which had consistently strong scores for the relevant PC across different analyses. If the anchor trait score on a given PC was negative, the entire component (both trait scores and window loadings) was multiplied by -1. This convention was applied consistently for all analyses.

### UMAP visualisation

We used UMAP embedding of the 403 traits using the uwot R package (McInnes et al. 2018), applied to the same scaled 403-trait × 51,907-window matrix used for SVD.

### Null permutation analyses

To assess the dependence of the recovered latent structure on the directionality and local genomic organisation of effects, we generated two different types of permuted null datasets, each with 100 replicates. First, entry-level sign randomisation: each entry of the matrix was independently sign-randomised, serving as a global null (Fig. 1c) and baseline for Procrustes similarity comparisons (Suppl. fig. 12). Second, within-block permutation: window effect values were shuffled within independently defined LD blocks (Berisa and Pickrell 2016); or within 500 kb or 250 kb blocks (Suppl. figs. 12, 14, and 21), preserving broad-scale genomic structure while disrupting locally coherent directional signal.

### Procrustes similarity

To compare latent trait configurations across analysis conditions, we computed the normalised Procrustes similarity (Gower 1975; Gower and Dijksterhuis 2004) between pairs of trait-score matrices from the singular values of the cross product of the two trait score matrices. To ensure this was an interpretable null baseline, we also computed Procrustes similarity between the observed configuration and entry-sign-randomised matrices. Because Procrustes alignment maximises similarity after rotation, unrelated trait configurations will usually retain non-zero raw similarity. We therefore report baseline-adjusted Procrustes similarity, which was rescaled so that the expected (mean) sign-randomised similarity was 0 and perfect alignment was 1. Specifically, for raw similarity s, we calculated s’ = (s - E[s_null_])/(1 - E[s_null_]), where E[s_null_] is the expected (mean) similarity to sign-randomised matrices for the same number of components. Similarity values were computed for k = 4, 8, 25, 50, and 75 components.

### Phenotype category enrichment

We computed the Spearman correlation between the vector of window median effects for each trait and each PC loading vector across the 51,907 windows. We then calculated the median absolute Spearman’s rho within each phenotype category, as the directionality of correlation is arbitrary due to trait encoding. To assess whether the observed category-specific correlations exceeded those expected by chance, we performed 1,000 label permutations, in which phenotype-category labels were shuffled across all 403 traits simultaneously, and the mean rank of the absolute Spearman’s rho per permuted category per PC was recomputed. Significance was determined by comparing the observed mean rank absolute rho to the resulting null distribution, with p-values estimated as the proportion of permutations exceeding the observed value.

### Removal of GWAS-significant loci

To test whether the recovered latent structure depended on established GWAS loci, we identified all windows containing at least one SNP with -log10(p) exceeding 7.301 (corresponding to p < 5e-8) in any of the 403 studies. We then extended each such window by a flanking buffer on each side (100 kb and 150 kb were tested; results here use 100 kb unless otherwise noted) to minimize residual LD, and removed all windows overlapping the resulting merged intervals. This reduced the matrix from 51,907 to 12,688 windows in the case of 100 kb flanks. SVD was reapplied to the retained windows using identical settings to that used in the full dataset SVD. Category enrichment analyses were repeated on the GWAS-depleted decomposition (Suppl. fig. 20).

### Proximity to removed loci

To test whether the latent structure recovered from GWAS-depleted windows might reflect residual LD contamination from removed loci, we computed the distance from each of the top-loaded windows (top 1%, 2%, and 5% by absolute loading for each PC) to the nearest removed GWAS window. Adjacent top-loaded windows were collapsed to a single representative (the central window of each contiguous run) before computing distances. A chromosome-matched null distribution was constructed by sampling an equal number of non-adjacent windows per chromosome from the retained set, repeating this procedure 2,000 times. A one-sided permutation test compared the area between the observed and mean null cumulative distribution functions.

### Genomic localisation of leading windows

We characterised the genomic context of top-loaded windows using two complementary approaches. First, we computed the distance from each window to the nearest protein-coding gene body, using Gencode v19 annotations (Frankish et al. 2021) and GenomicRanges R package (Lawrence et al. 2013). Windows directly overlapping a gene body were assigned distance zero. Distances were binned as: genic (0 bp); 0-50 kb; 50-100 kb; 100-250 kb; > 250 kb. For each PC and each top-loading fraction (top 1%, 2%, 5%), we calculated the log2 ratio of the observed fraction in each bin to the mean fraction across 1,000 chromosome-matched null replicates. Significance was assessed empirically from the null replicate distribution.

Second, we assigned each 50 kb window to a mutually exclusive functional annotation class using Gencode v19 GTF annotation: promoter (within ±2 kb of any annotated transcription start site), exonic (overlapping an exon but not a promoter), intronic (overlapping a gene body but not an exon or promoter), and intergenic (no overlap with any gene body or promoter). For each PC and annotation class, we computed the cumulative ratio of observed to genome-wide background overlap fraction, ranked from the top 1% of loaded windows to all windows, with 95% confidence intervals derived from 100 chromosome-matched null replicates.

### Gene-set enrichment analysis

To characterise the biological processes associated with strongly loaded genomic windows, each window was assigned to its nearest protein-coding gene using Gencode v19 (Frankish et al. 2021), regardless of distance. Adjacent top-loaded windows were collapsed before assignment by retaining the central window of each contiguous run. Gene-set enrichment was performed using clusterProfiler (Wu et al. 2021). enrichGO() was used to test for enriched GO terms (Biological Process, Molecular Function, Cellular Component); MSigDB Hallmark gene sets and curated canonical pathways were tested with enricher(), with gene sets retrieved via the msigdbr R package (Liberzon et al. 2015; Subramanian et al. 2005). All tests used a background gene set comprising the nearest-gene annotations for the complete set of retained windows, Benjamini–Hochberg FDR correction, and FDR-adjusted thresholds of p < 0.05 and q < 0.2. Enrichment was computed for the top 2% of windows by absolute loading for each PC. For visualisation, only GO terms with an intersection size of at least 15 genes are shown. Redundant GO terms were removed using GOSemSim (Yu et al. 2010): terms were sorted by adjusted p-value and any term with Wang semantic similarity ≥ 0.7 to a higher-ranked retained term was discarded (Wang et al. 2007).

## Author contributions

OKS conceived the research and analysed all data. LLMs were used to assist in the coding and data analysis and to check for consistencies in manuscript grammar and format. The code for performing the analyses and producing all figures is available at https://github.com/osilander/distal-polygenic-architecture.

## Acknowledgements

Thank you to J. O’Sullivan, S. Gokuladhas, C. Miller, and T. Portlock for discussions during the development of this work. This work was funded by a gift from John Werry through the University of Auckland foundation.

## Supplementary Figures

**Supplementary figure 1.**
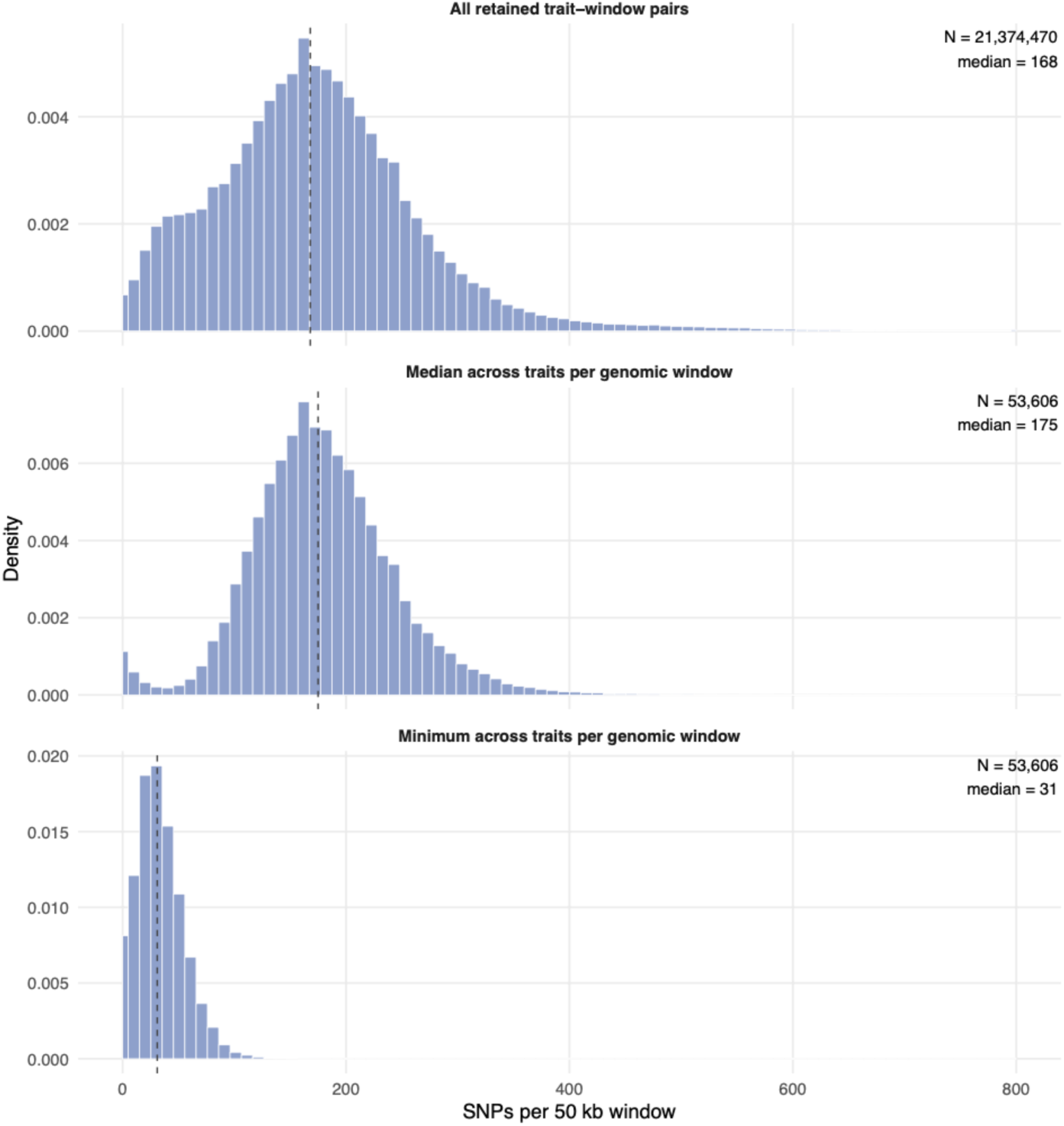
Distribution of SNP counts per 50 kb window across traits. Histograms show the number of SNPs retained in each window after quality filtering, stratified by trait. Top panel: all windows across all 403 traits. Middle panel: median number of SNPs within each window across all traits. Bottom panel: Minimum number of the SNPs within each window across all traits. The trough in the middle panel occurs at approximately 30 SNPs, which was used as a threshold cutoff for all analyses. Windows with fewer than the minimum SNP threshold were excluded from all downstream analyses.

**Supplementary figure 2.**
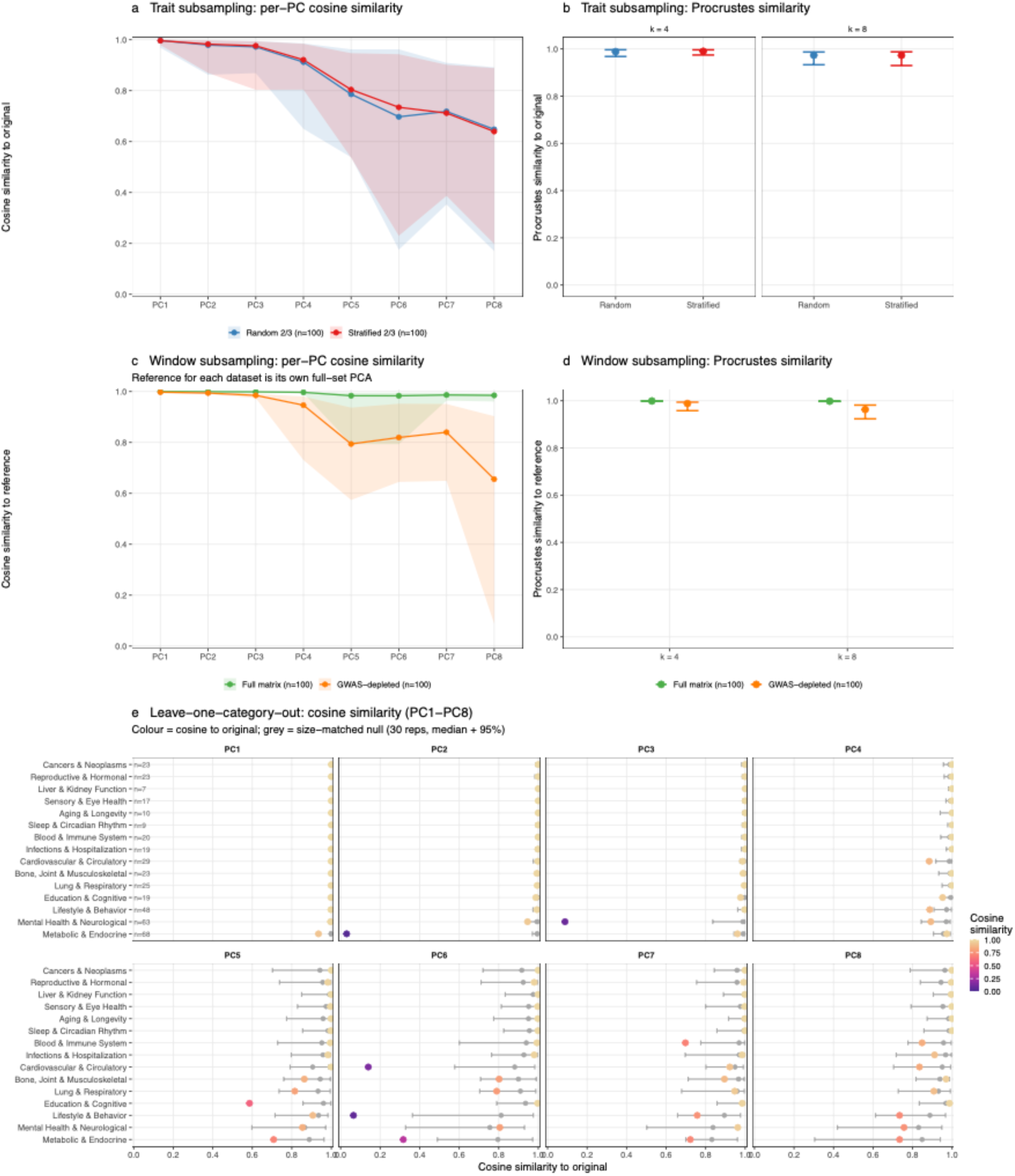
Robustness of decompositions. a. Median cosine similarity between each corresponding PC score vectors after subsampling 2/3 of all traits, and the PC from the full decomposition, across 100 random and 100 trait-group stratified replicates. Ribbons show the 2.5–97.5% intervals. PC similarities are extremely high until PC4, with a slow decline after. b. Normalised Procrustes similarity between the subsampled and full latent trait configurations for the first 4 (left panel) and 8 (right panel) PCs; error bars show the 2.5–97.5% interval. As with a., trait space is robust to trait removal. c. and d. Analogous results as those shown in a. and b., but for 2/3 genomic window subsampling of the full and GWAS-depleted matrices. In both cases, window subsampling leaves the first four PCs almost identical to their respective full data sets. e. Leave-one-trait-category-out cosine similarity for PC1–PC8. Each trait category was removed in turn and the decomposition rerun on the remaining traits; the resulting components were matched to their full-decomposition counterparts by greedy maximum-cosine assignment, and the per-PC cosine similarities are shown. Low cosine similarity indicates that the category is structurally important for that axis. Grey bars show the median and 95% interval of a size-matched null, in which an equal number of randomly drawn traits as those in the trait group were removed (30 replicates). Categories are ordered by PC1 cosine similarity. The sensitivity of PCs to trait-category removal corresponds well with the correlations of PCs and trait categories (Fig. 1d; Suppl. fig. 3). For example, PC1 is robust to the removal of almost any trait group; all later PCs are heavily disrupted with the removal of one or more groups. For example, PC3 is disrupted by removal of Mental Health and Neurological traits; PC4 by removal of Cardiovascular and Lifestyle trait groups; and PC7 by removal of Blood and Immune traits. Note that later PCs explain similar and smaller amounts of variation and can thus interchange with one another; a low cosine for later components may be due to reordering among components rather than genuine instability.

**Supplementary figure 3.**
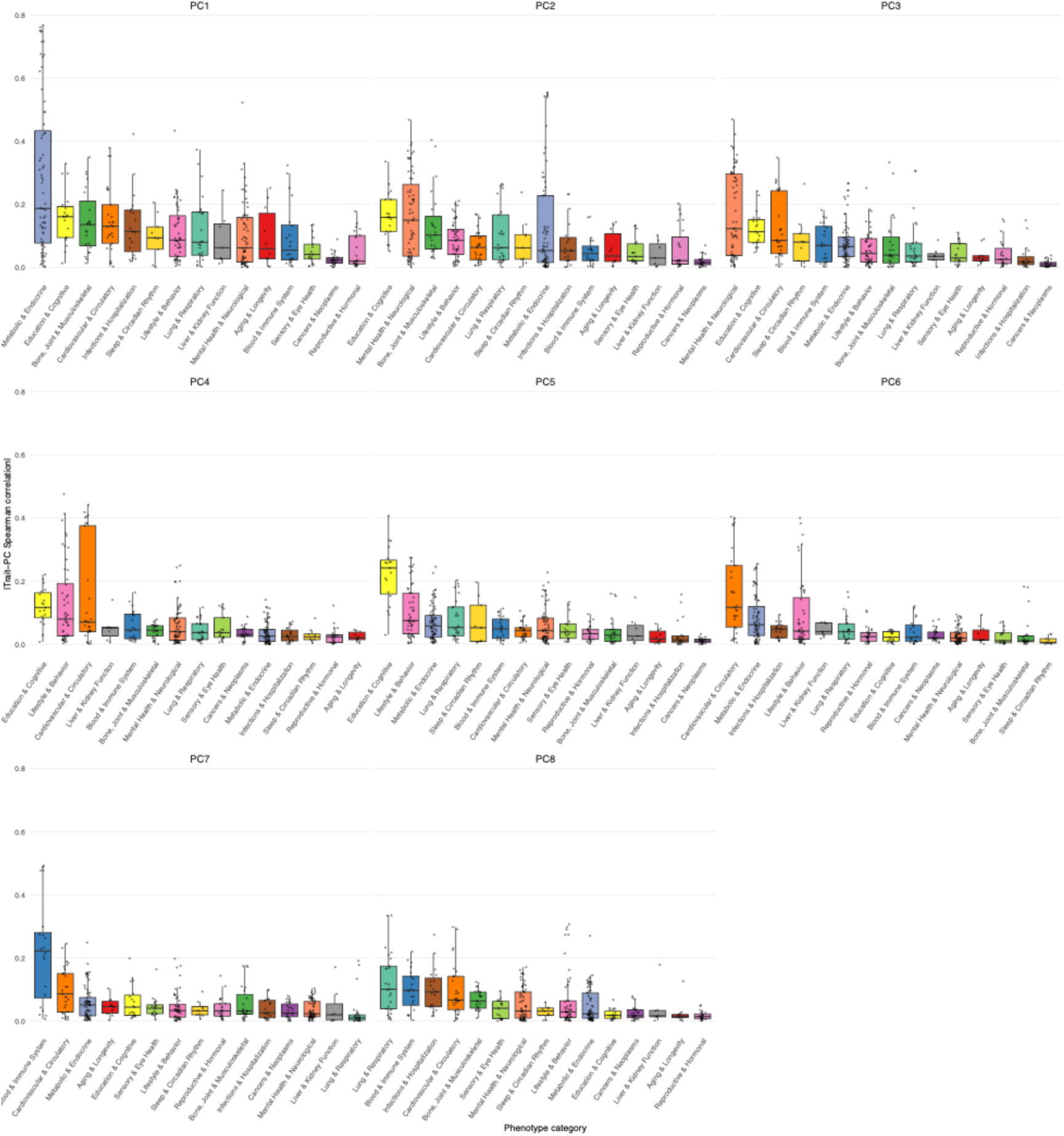
Absolute Spearman correlation (rho) between window PC loadings and window median effect sizes for each individual trait, separated by trait category, for PCs 1-8. Each point indicates an individual trait; boxplots indicate the median and upper and lower quartiles within categories. Some PCs are generally more correlated with traits in a single category. For example, PC7 window loadings are well correlated with window median effects in many Blood and Immune cell traits, but poorly with traits in most other categories. Conversely, window median effects of Metabolic and Endocrine traits are well correlated with PC1 loadings, but poorly with all other PC loadings. Absolute values are used for correlations, as the directionality is arbitrary due to trait encodings and PC orientation. This analysis provides convergent evidence for the Leave-one-trait-category-out analysis in Suppl. fig. 2.

**Supplementary figure 4.**
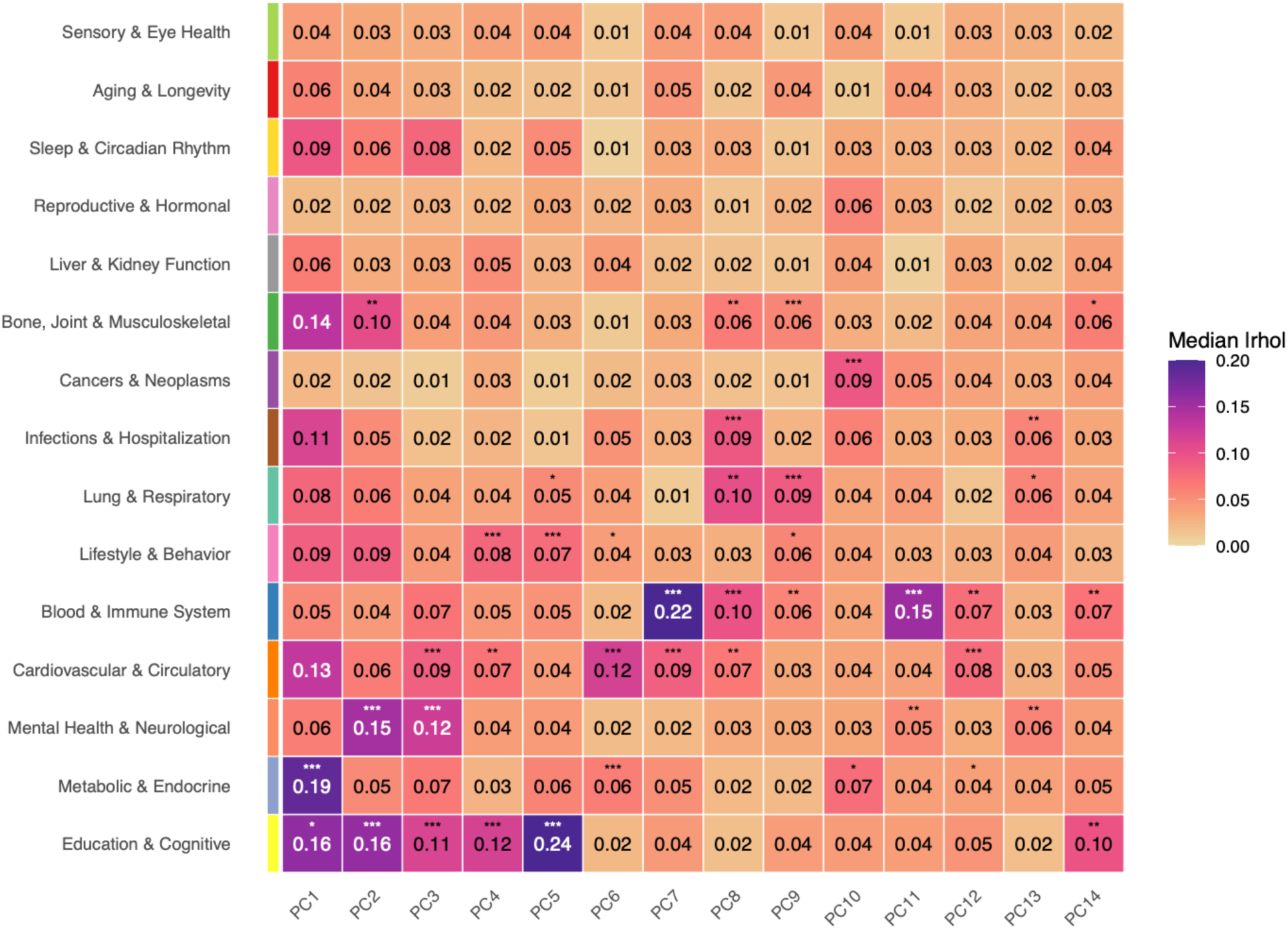
Median absolute Spearman correlations (rho) between window PC loadings and window median effects for all traits in each category (see Supplementary figure 3). This is the same data as that shown in figure 1d but annotated here with the Spearman correlation values. Some trait groups correlate well with only a single PC (e.g. Cancer and Neoplasms with PC10); others with multiple (e.g. Education and Cognitive traits). Conversely, some PC loadings correlate well with only a single trait group (PC7 with Blood and Immune); others correlate with several (PC1). Significance values are derived from 1,000 label permutations in which trait category labels are swapped and the per trait median values recalculated. The colour scale is capped at 0.2.

**Supplementary figure 5.**
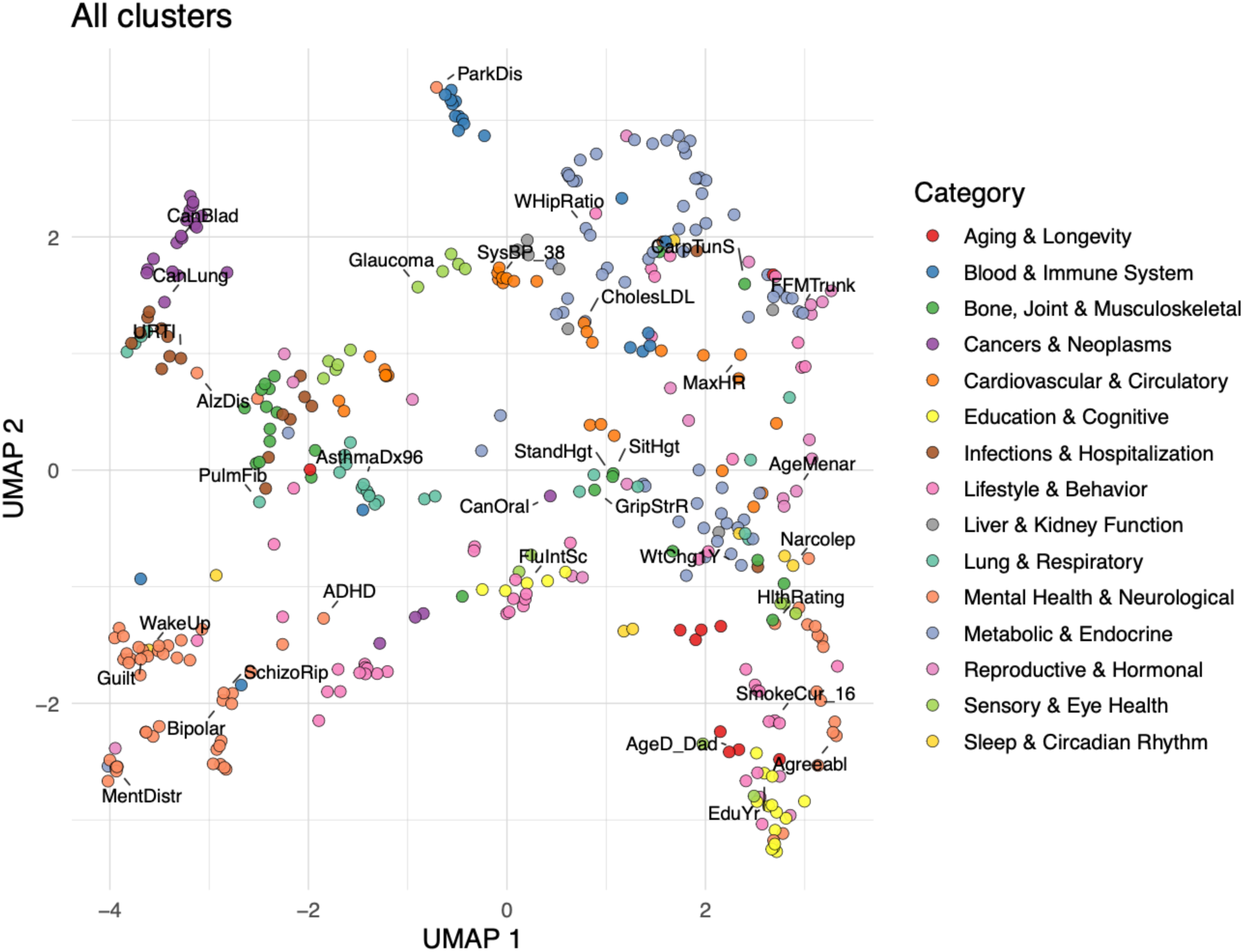
UMAP projection of traits based on their genome-wide window effect. Clustering reflects similarity in the genome-wide pattern of window median effects. A subset of traits are labelled for orientation.

**Supplementary figure 6.**
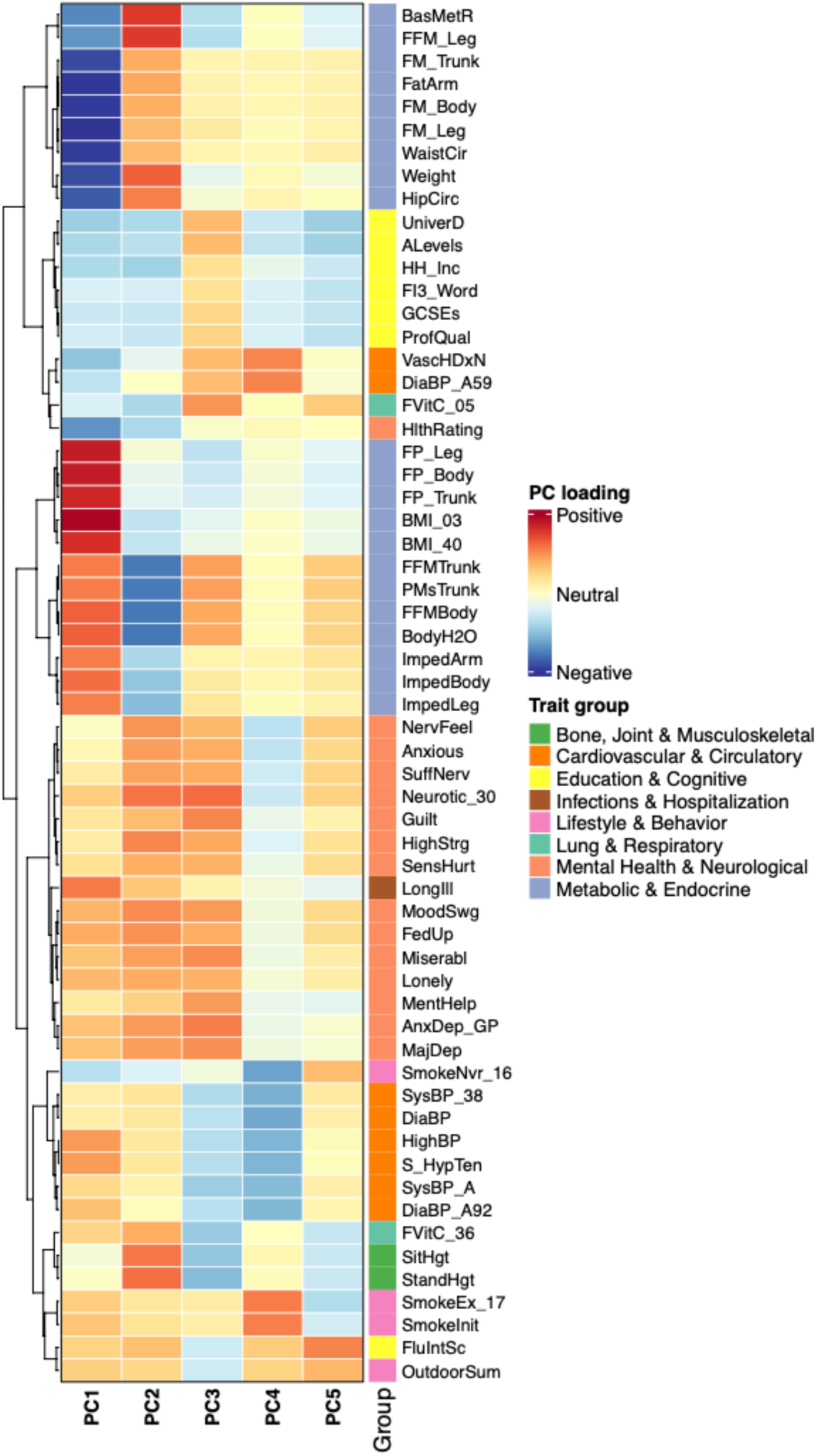
PC loading heatmap for the 60 traits with the largest total absolute loadings (sum) across PCs 1-5. Rows are clustered by Ward’s method and coloured on the right side by trait category. Most traits cluster as expected, although in some cases trait encodings can result in similar traits having directly opposed loadings or opposite traits having similar loadings (e.g. fat percentage trunk (FP_Trunk) and fat-free mass trunk (FFMTrunk)).

**Supplementary figure 7.**
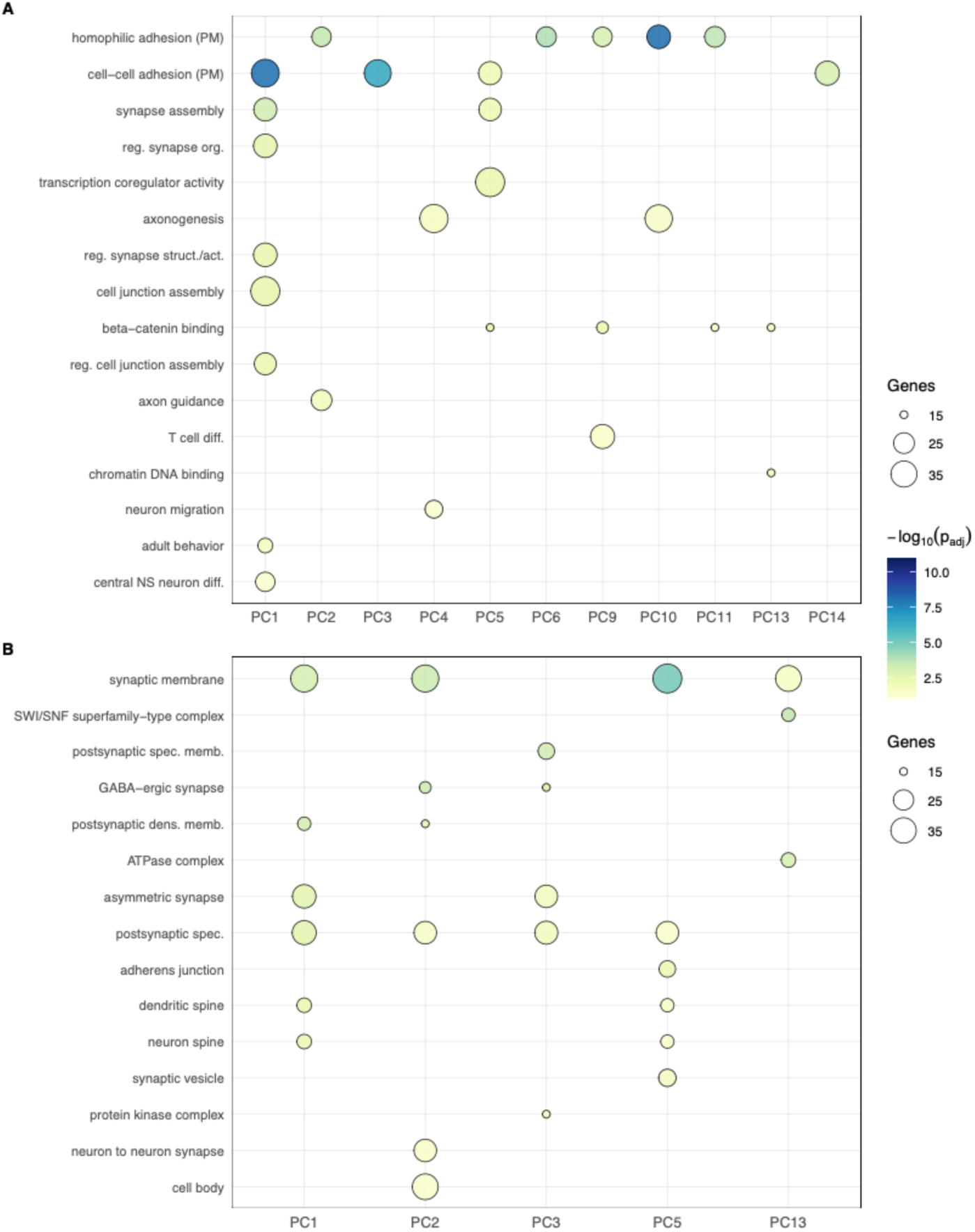
Gene Ontology enrichment for windows in the top 2% (1,038 windows) of absolute loadings for PCs 1-14, using the most proximal gene for ontology annotation. Only PCs with at least one category having significant enrichment for either Biological Process (BP) or Cellular Component (CC) are shown (for example, no significant enrichment was observed for PC7 or PC8 in either BP or CC so neither PC7 nor PC8 are shown). Note that the sets of categories are not mutually exclusive. While the vast majority of enrichment categories relate to neuronal function, PC9 shows enrichment for genes functioning in T cell differentiation. This latent axis is most correlated with traits in the Blood & Immune System and Lung & Respiratory trait groupings.

**Supplementary figure 8.**
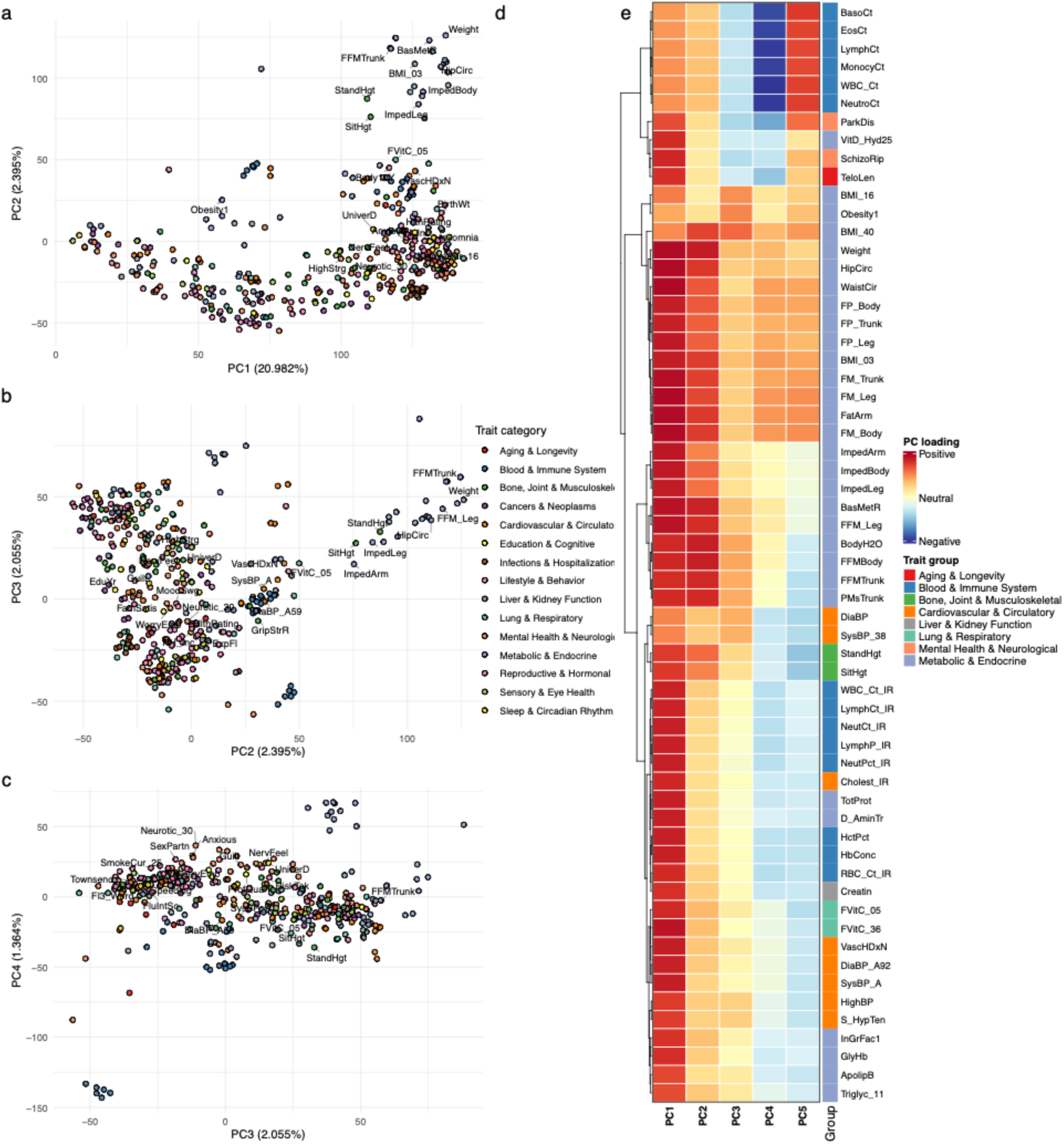
PCA of absolute GWAS effect sizes (betas) aggregated into 50 kb windows. Trait projections of the first few PCs exhibit less coherence than signed values; traits grouped by PC loadings are also less coherent (e.g. grouping telomere length and schizophrenia; see Figure S5).

**Supplementary figure 9.**
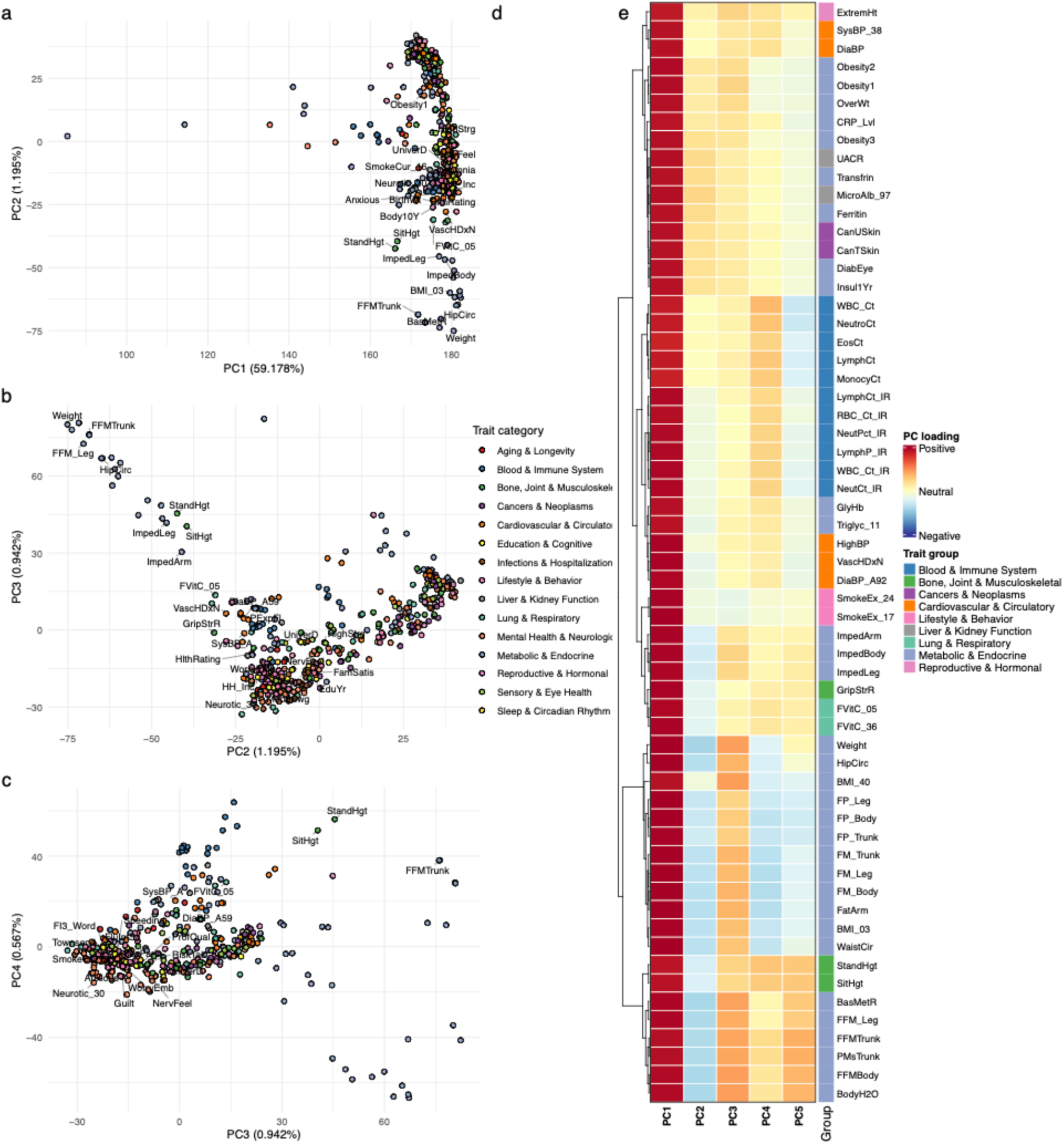
PCA of absolute (unsigned) median effect sizes across 50 kb windows. As with absolute value of individual betas, trait projections of the first few PCs exhibit less coherence than signed values; and trait groups by PC loadings are also less coherent (e.g. grouping diabetes-related eye conditions and skin cancer; see Figure S5).

**Supplementary figure 10.**
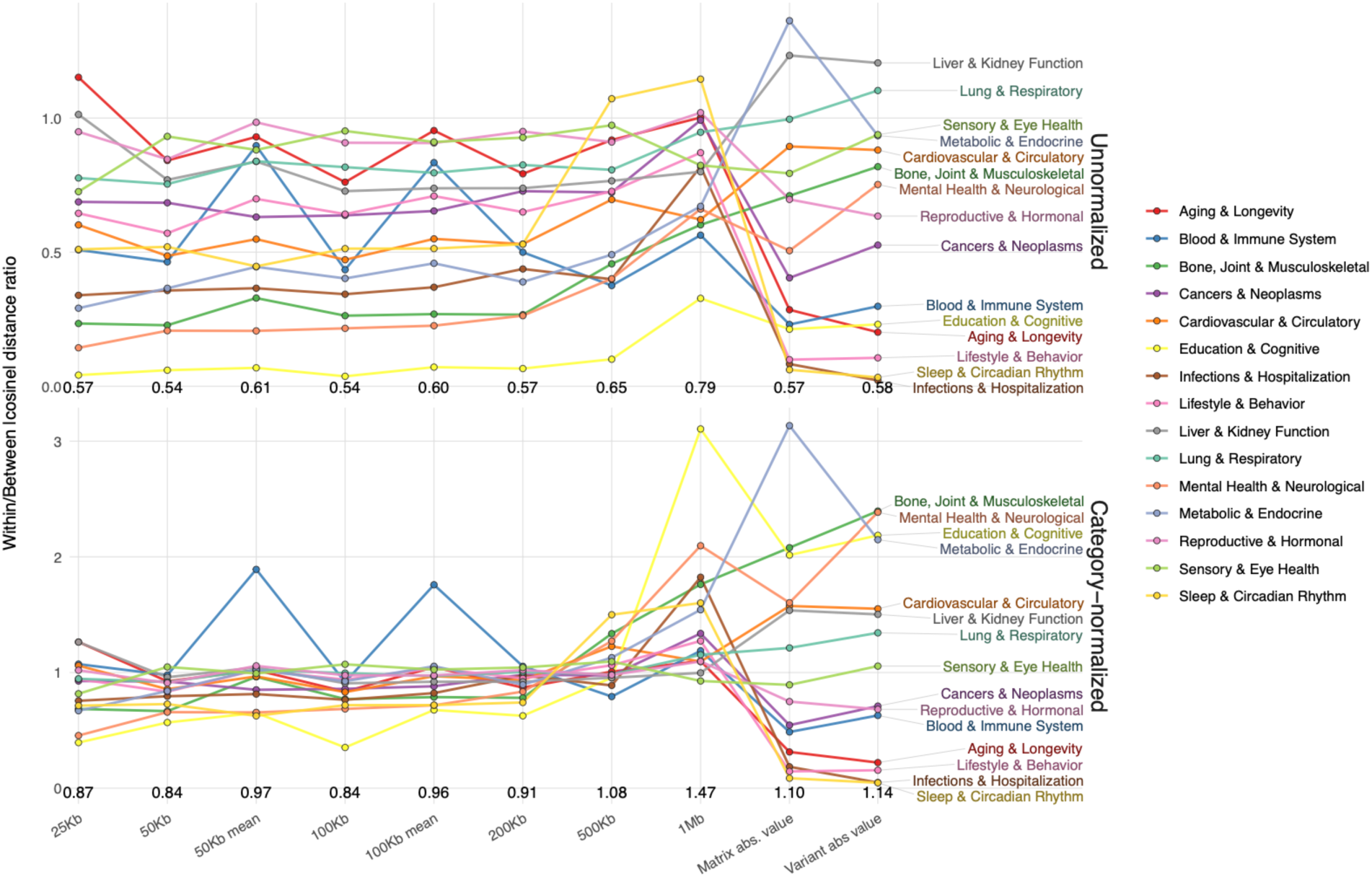
Category coherence in PC 1-4 space, across genomic window treatments. For each trait, absolute cosine similarities to all other traits are computed, split into within- and between-category medians, converted to cosine distances (1 – abs(cosine)), and summarized as the ratio of within-distance to between-distance for each phenotype category. Ratios below 1 indicate that traits cluster more tightly with their own category than with others. Top: raw category ratios, indicating that some trait categories are inherently more coherent, such as Education and Cognitive traits or Cardiovascular and Circulatory traits. Bottom: category ratios, normalized by each category’s mean across the six base window sizes (25 kb – 1 Mb) to highlight relative changes across analysis conditions. For each analysis condition, the mean across categories is annotated. The analysis conditions are: median signed effects across six different window sizes (25 kb – 1 Mb); mean signed effects at 50 kb and 100 kb; median absolute value effects, and the absolute value of median signed effects. The 50 kb and 100 kb windows using median values show the clearest and most consistent coherence and are used for all presented results.

**Supplementary figure 11.**
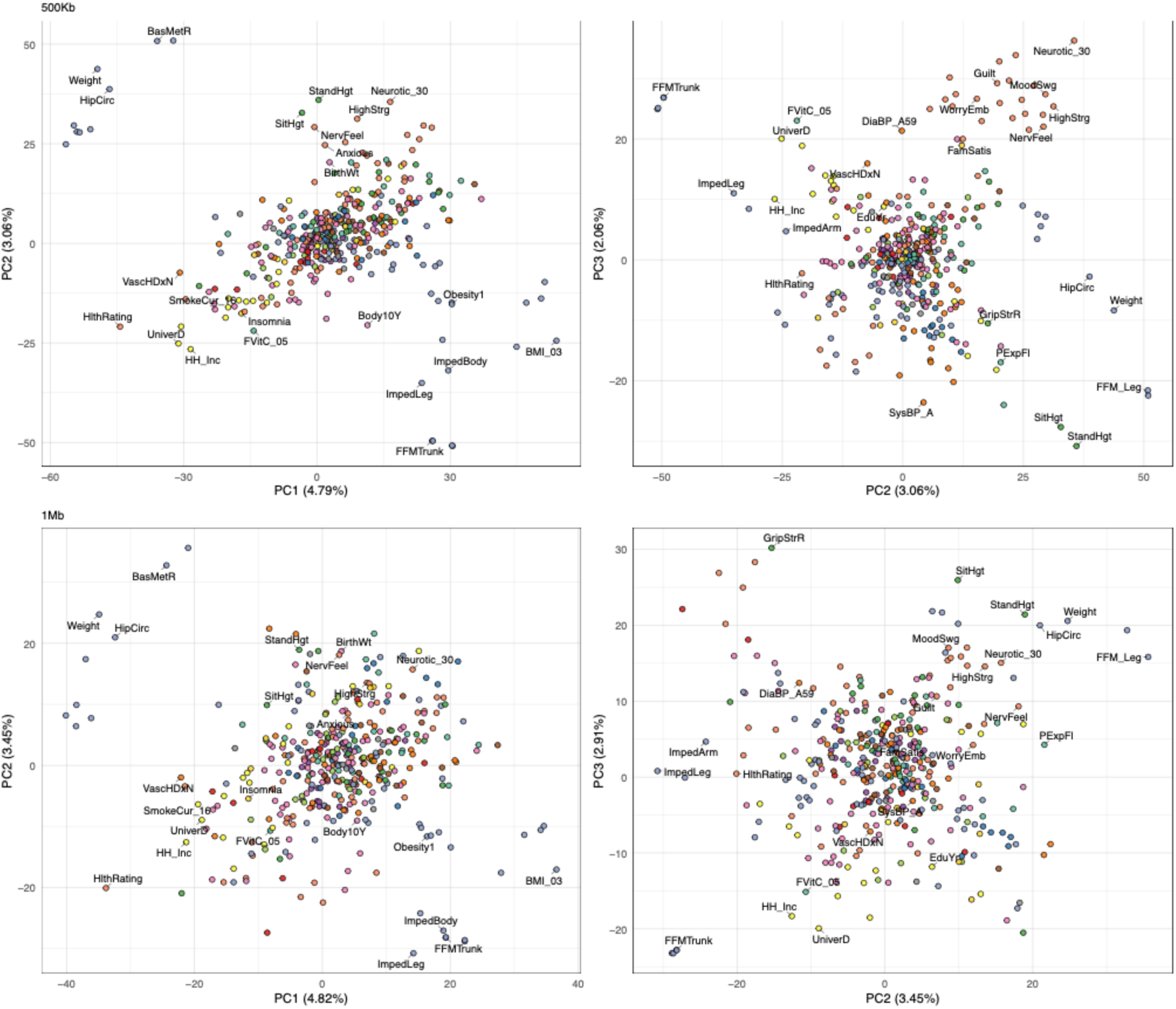
Trait biplots for median effects calculated across 500 kb (top row) and 1 Mb (bottom row) windows. Traits are coloured by category, with selected traits labelled for orientation. To simplify comparisons, PCs are oriented such that they match PC-specific anchor traits from the 50 kb window analysis. Orientation occurs in the following manner: for each PC, the score of the anchor trait is determined. If the value is the opposite sign of the trait in the 50 kb analysis, the entire PC (both trait scores and window loadings) is multiplied by −1. The anchor traits are: BMI_03 for PC1; standing height for PC2; and Neurotic30 for PC3. Trait category coherence is clearly lost for PC3 at the 1 mb window size.

**Supplementary figure 12.**
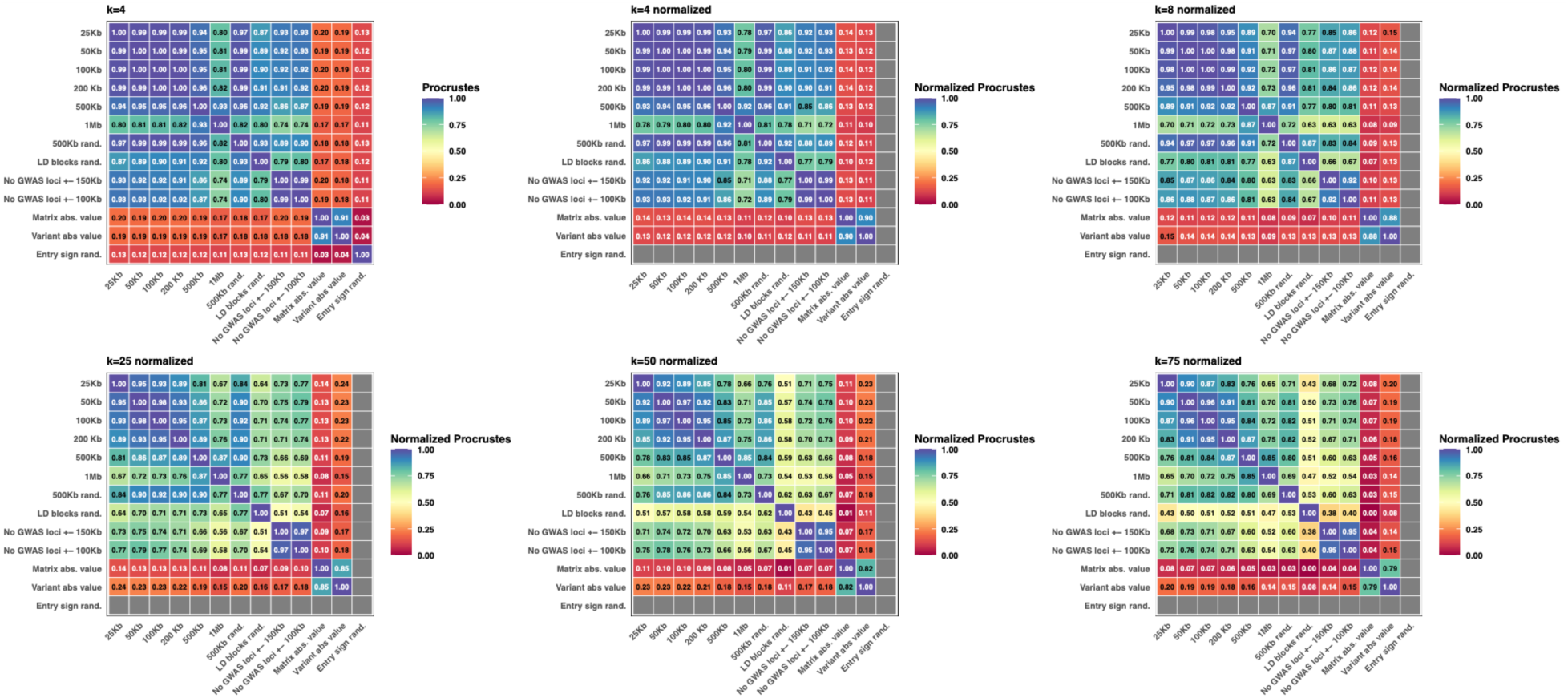
All-by-all Procrustes similarity between SV decompositions for 12 window block analyses or window permutations. Within-block permutations are performed by breaking the genome into 500 kb blocks, or by using blocks assigned by Berisa and Pickrell (Berisa and Pickrell 2016), randomizing window effects within blocks, and recalculating the SVD. Sign randomization is performed by multiplying each window (here, 50 kb) within each trait by -1 or +1, and acts as a baseline null in Procrustes similarity (Fig. 1c). GWAS-removed analyses are 50 kb windows, and include removal of both 100 kb and 150 kb flanks on each side. The top left heatmap includes the sign-randomized matrix, which exhibits minimal similarity to any other SVD. All later heatmaps indicate Procrustes similarity using the sign-randomised matrix values for rescaling such that perfect similarity is 1 and complete similarity is 0. From left to right and top-to-bottom, the Procrustes similarities are calculated using k = 4, 8, 25, 50, and 75, where k is the number of latent SVD components included.

**Supplementary figure 13.**
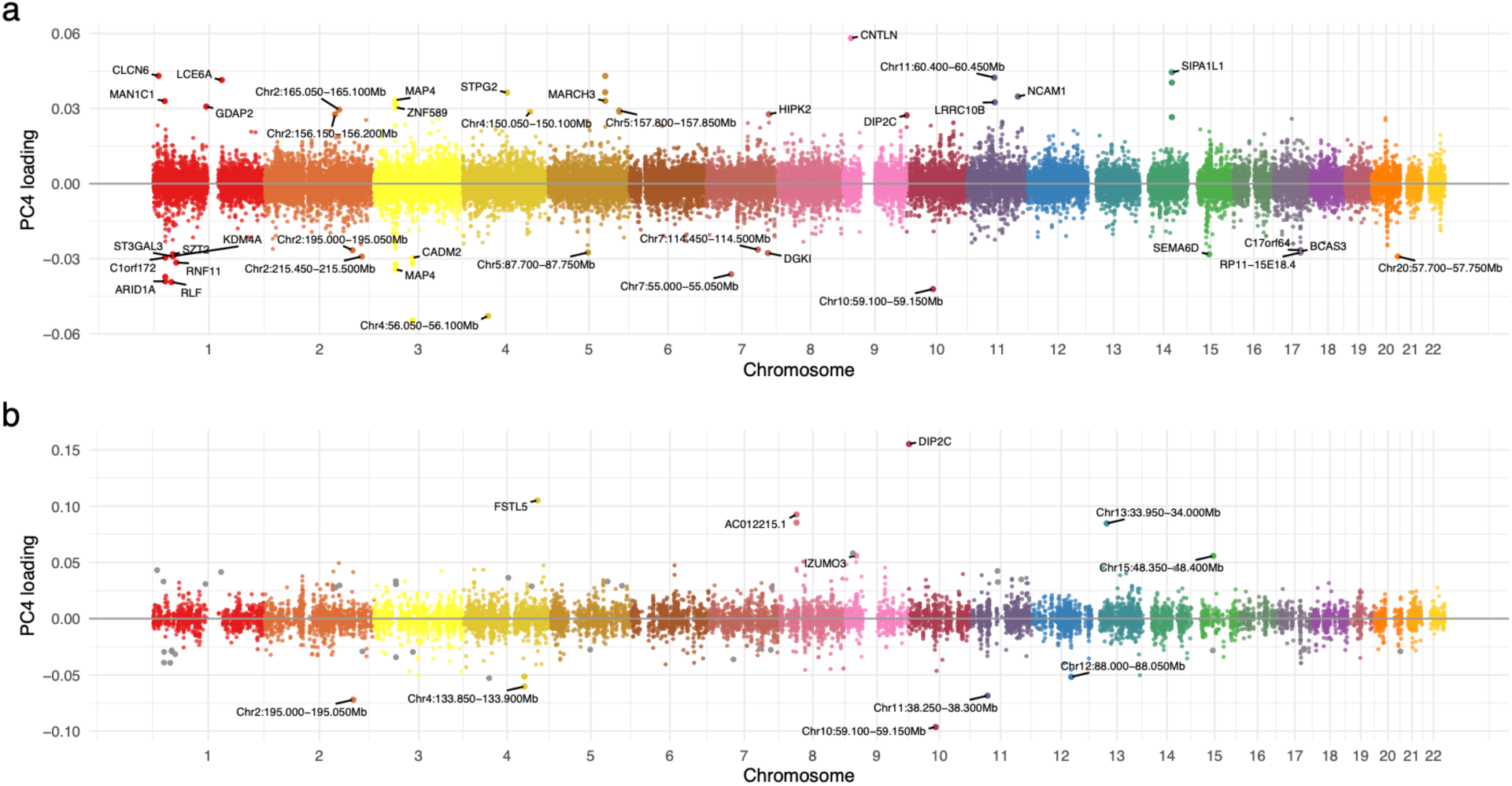
PC4 window loading profiles before (a) and after (b) removal of GWAS-significant windows and flanks (±100 kb). Points in the top 0.1% of absolute loadings are labelled with the name of the overlapping genic region; if no genic overlap is present, the genomic coordinates are indicated. Adjacent top-0.1% windows are merged so that only a single mid-region point is labelled.

**Supplementary figure 14.**
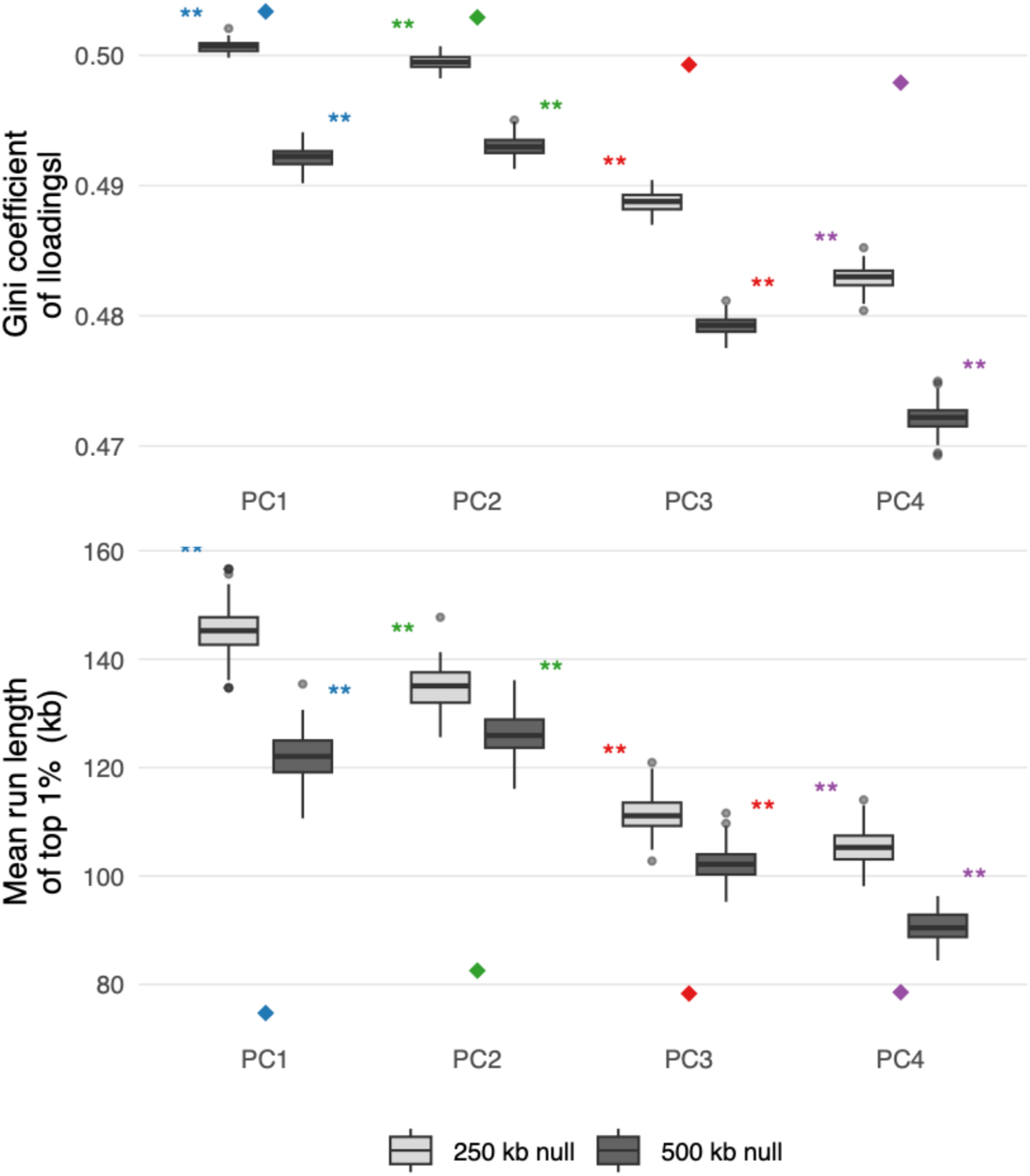
Concentration and spatial clustering of window loadings after PCA. Boxplots show the null distributions of the Gini coefficient of |loadings| (higher = more unequal) and the mean run length of top-1% windows in kb (shorter = more spatially clustered). The observed value for each PC is shown as a coloured diamond. Null distributions come from 100 within-block permutations, in which window effect values are shuffled within 250 kb or 500 kb genomic blocks independently for each trait, retaining broad-scale LD structure while disrupting locally concentrated signal. Asterisks indicate empirical significance (one-sided: upper tail for Gini, lower tail for run length; p < 0.01 in all cases). Results are shown for PC1 - PC4. PCs from each permutation are matched to the observed PC1 - PC4 by greedy maximum absolute cosine similarity. PC1 is matched first, then PC2 from the remaining unmatched permuted PCs, with sign correction applied such that each matched permuted PC points in the same direction as its corresponding observed PC.

**Supplementary figure 15.**
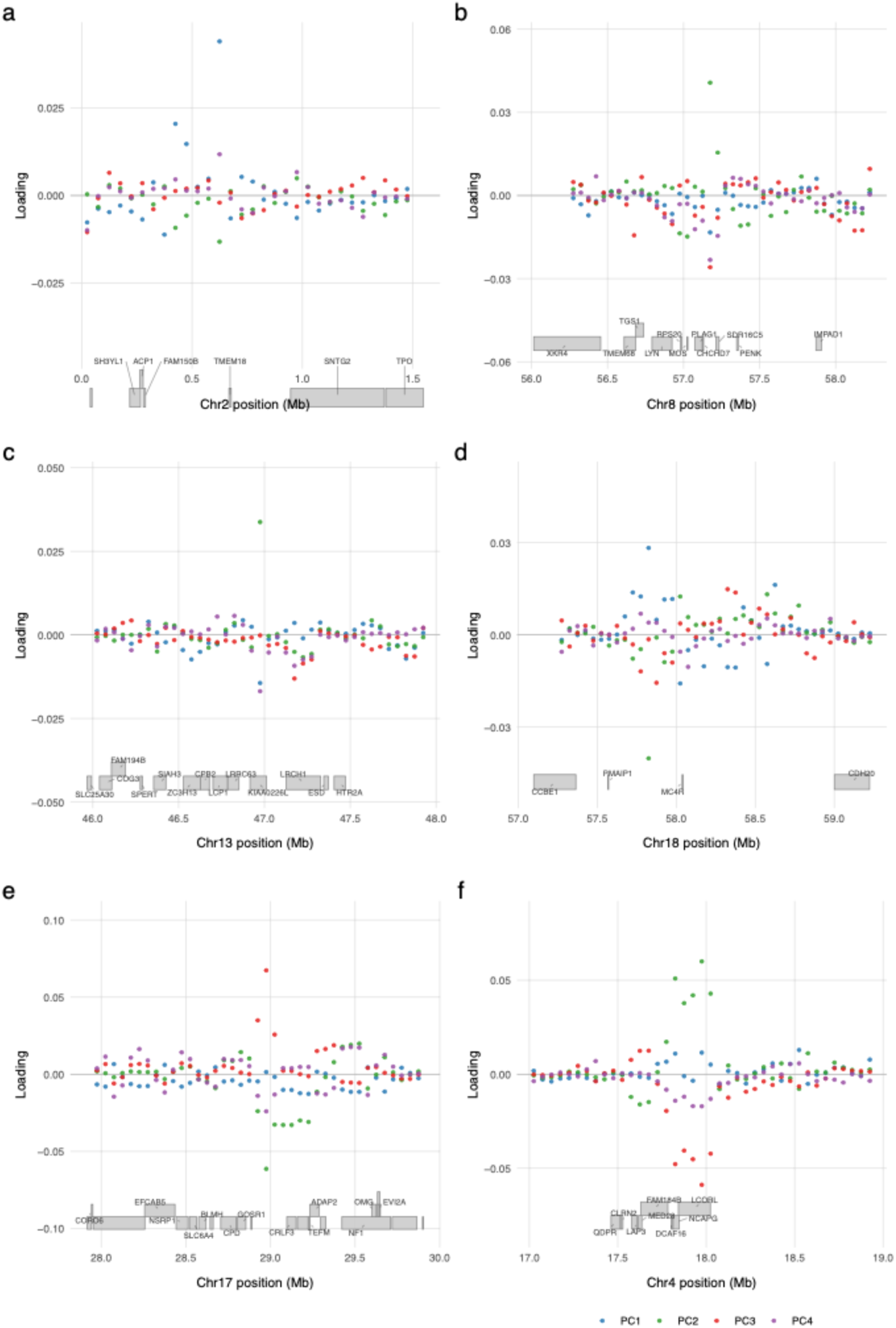
Zoomed regional plots for six regions with high PC loadings. Each panel shows the loadings on PC1 – 4 with the annotated genes for that region. In many cases, two or more PCs exhibit strong loadings, although the combinations change (e.g. in panel d PCs 1 and 2 exhibit strong loadings at the MC4R locus; in panels e and f, PCs 2 and 3 exhibit strong loadings). Note that the direction of the loading is arbitrary and is a function only of the specific PC rotation selected.

**Supplementary figure 16.**
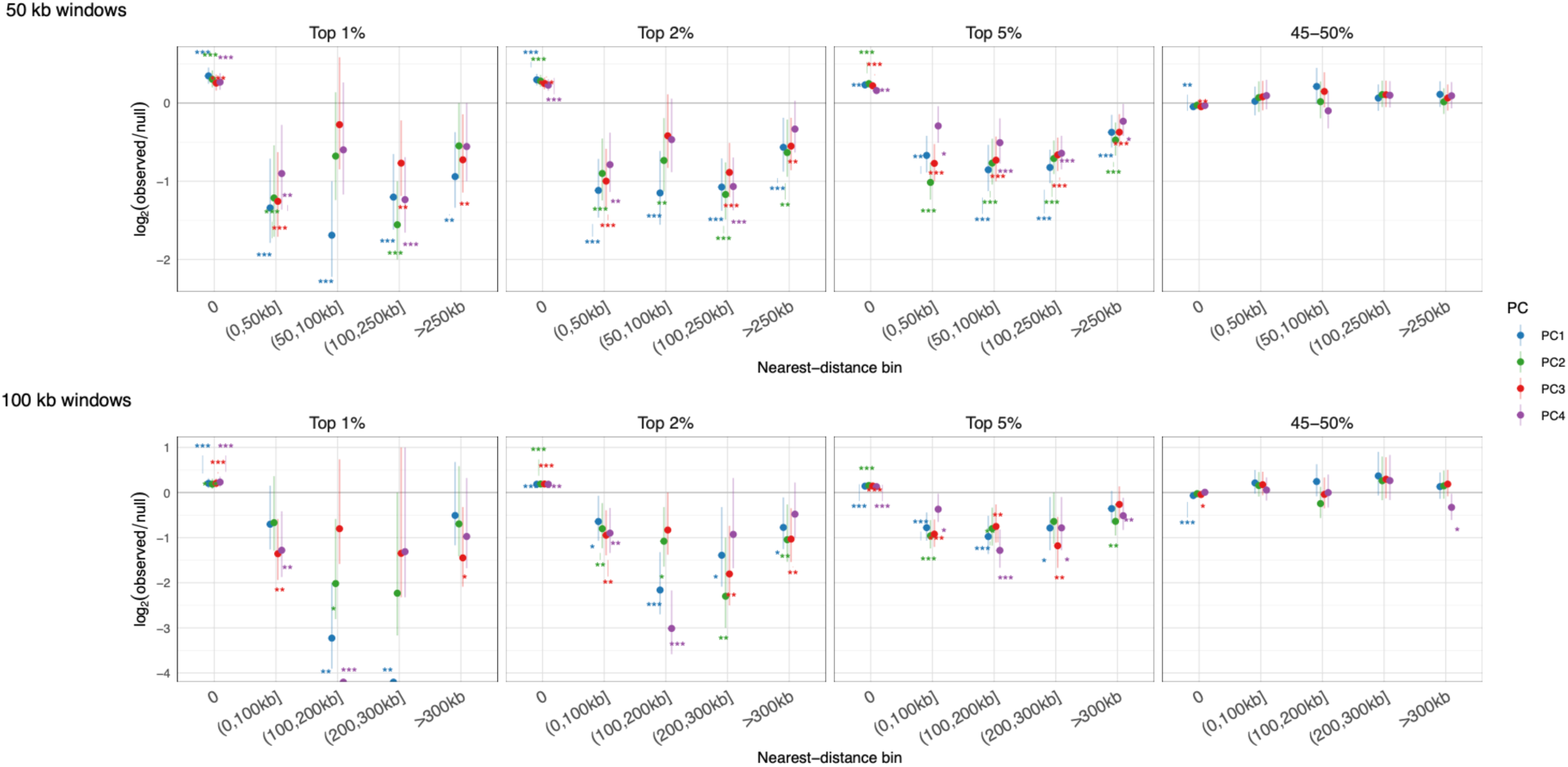
GWAS-locus enrichment for each distance bin (log₂ observed/null) for the top-loaded 1%, 2%, 5% of genomic windows, with the loadings at the 45–50% loaded windows indicated as permutation confirmation. The results for both 50 kb (top) and 100 kb (bottom) genomic windows are shown. In all cases, adjacent top-loaded windows are collapsed to a single representative (the middle window of each contiguous run) before computing distances. The null distribution is constructed by sampling an equal number of non-adjacent windows per chromosome from the same set, calculating the fraction in each bin for this null set, determining the ratio of the observed bin fractions to the randomised fractions, repeating this sampling 3,000 times. Whiskers indicate 95% confidence intervals, asterisks indicate significance (* p < 0.05; ** p < 0.01 *** p < 0.001).

**Supplementary figure 17.**
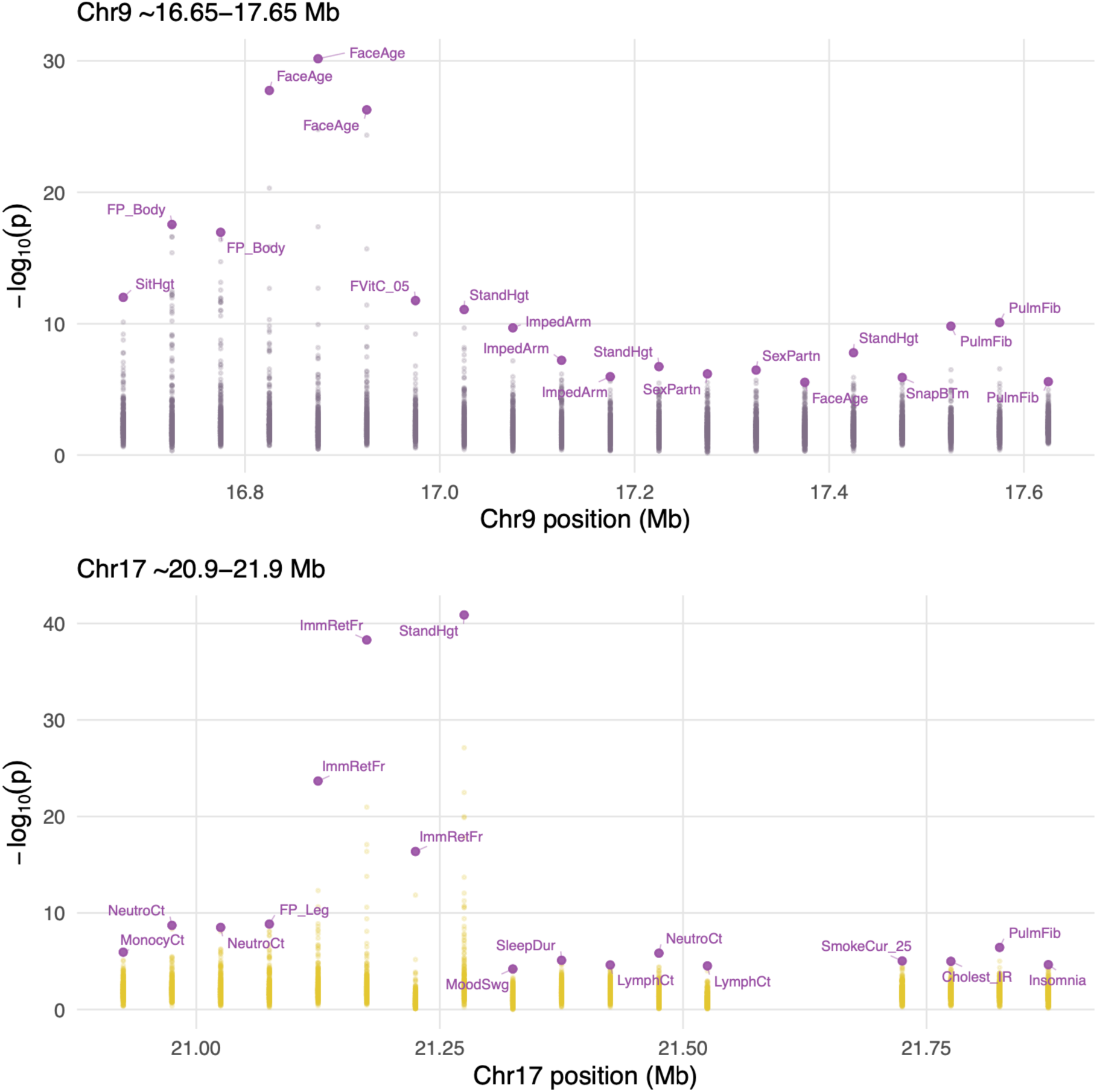
Genomic regions with strong PC loadings but no significant GWAS loci. Two loci are shown, the CNTLN/SH3GL2 locus at Chr9 17.2 Mb; and the C17orf51 locus at Chr17 21.45 Mb. Neither harbour GWAS-significant windows, but are heavily loaded for several PCs. The trait with the maximum -log10 p-value is indicated for each window. Four windows on the Chr17 locus had too few SNPs to be considered.

**Supplementary figure 18.**
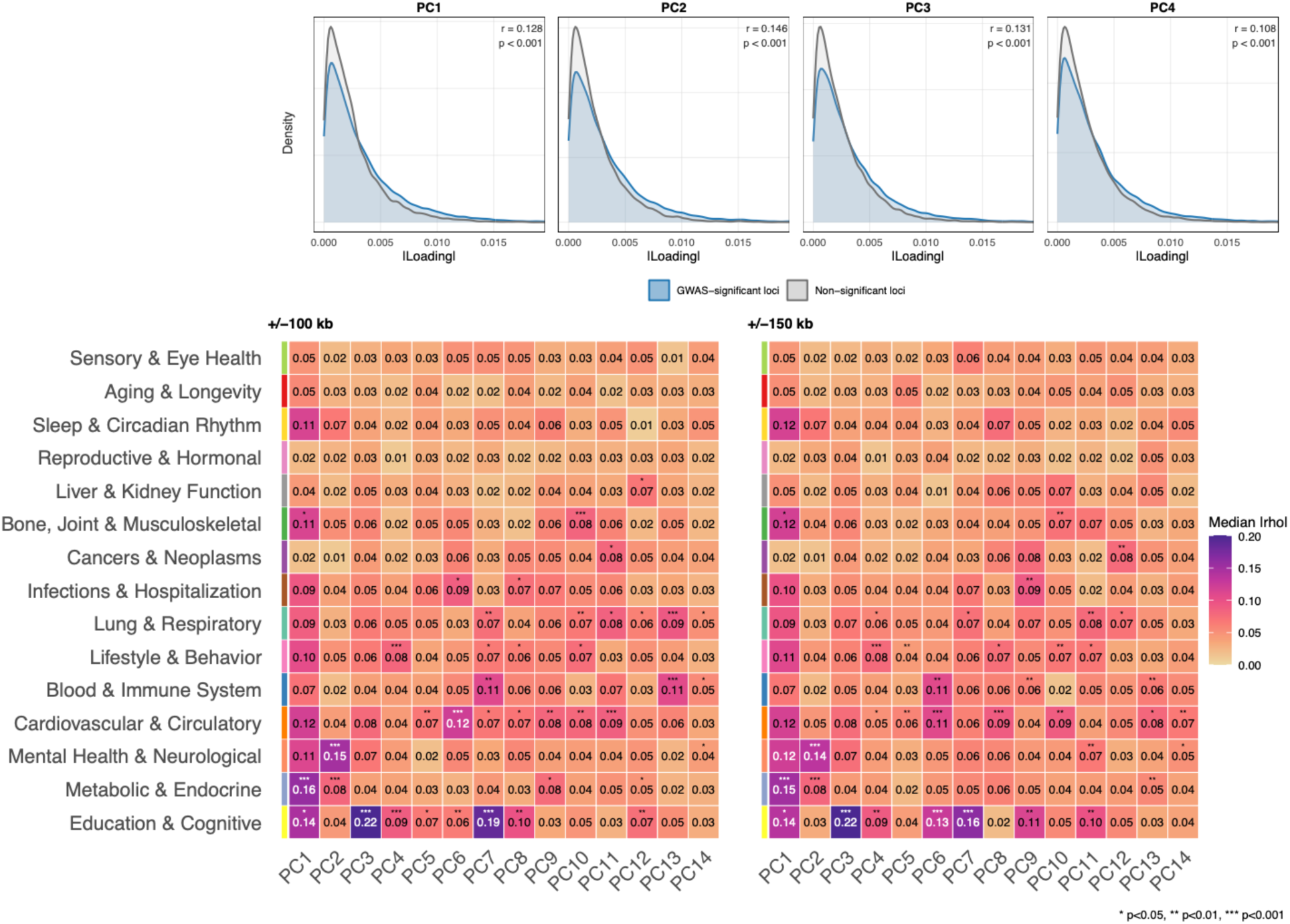
Upper panel: density plots comparing absolute loadings for PC1 – PC4 for GWAS-significant windows (blue) vs non-GWAS significant windows (grey; GWAS flanking regions are excluded from both groups). P-values indicate the results of a Wilcoxon test. In all cases, GWAS-significant windows exhibit far stronger loadings (p < 0.001); this corroborates the results shown in Suppl. fig 16, but considering all loci and without explicit distance-based statistics. Lower panel: Median absolute Spearman correlations for the median window effects vs. PC loadings for all traits in each category for PC1 – PC14. 1,000 trait label-permutations were used to infer significance (* p < 0.05; ** p < 0.01 *** p < 0.001). Results for GWAS loci ±100 kb flank removal are shown on the left; ±150 kb flank removal is on the right.

**Supplementary figure 19.**
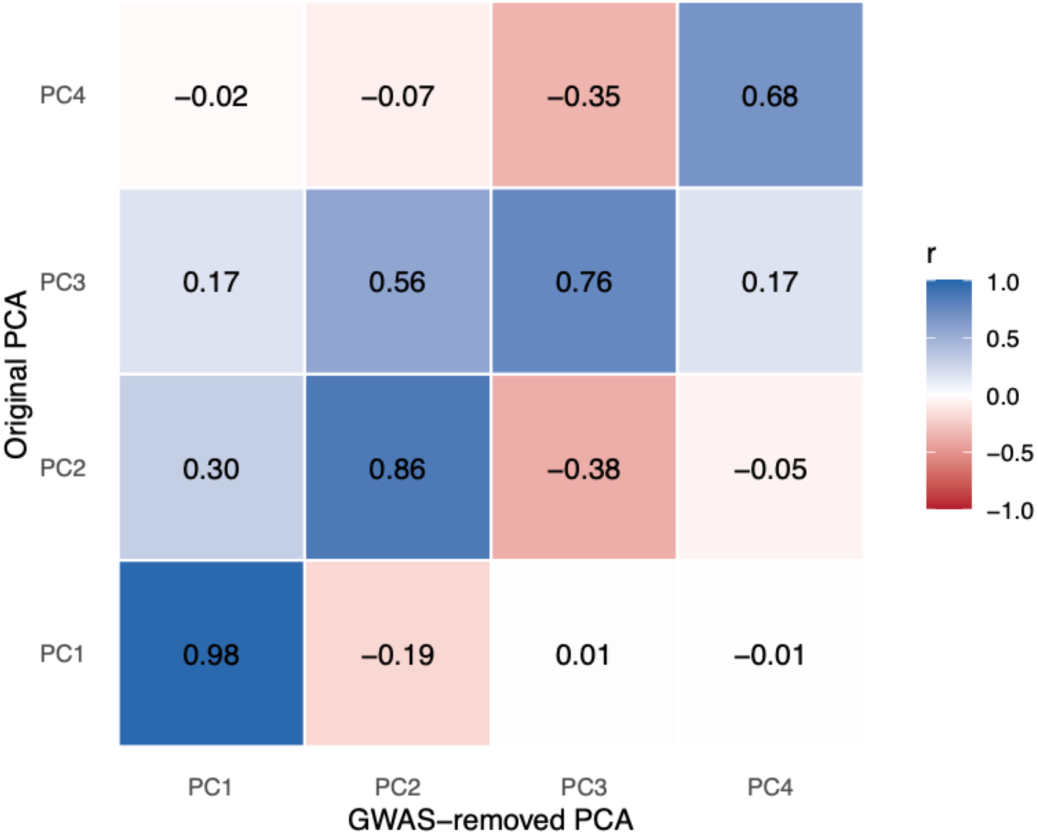
Pairwise correlations for PC1 – PC4 for the full dataset and the GWAS-removed data (±100 kb flank removal). Each GWAS-removed PC was sign aligned to the best-matching original PC, as determined via cosine similarity.

**Supplementary figure 20.**
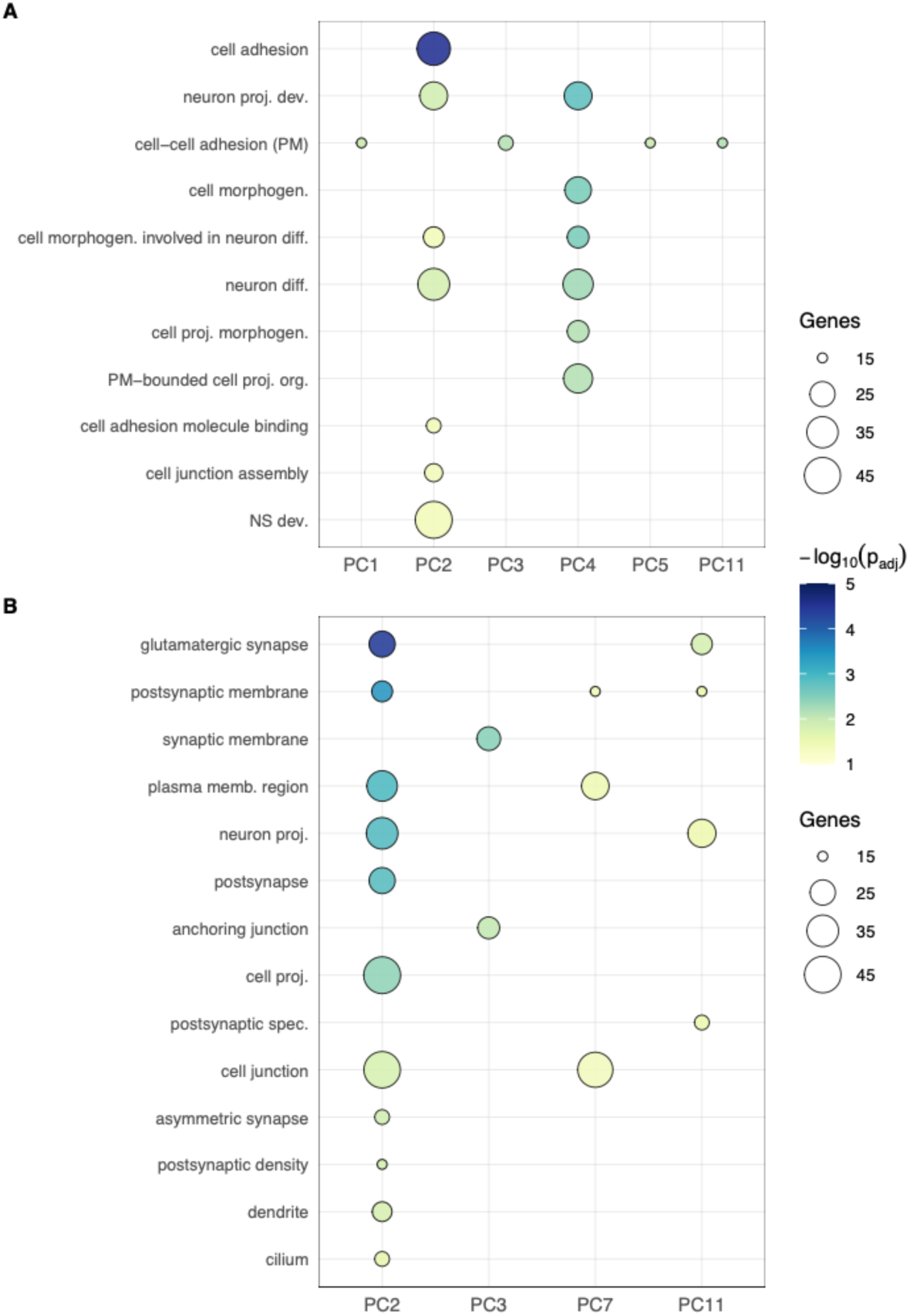
Gene Ontology enrichment for windows in the top 2% of absolute loadings (254 windows) for PC1 – PC14 of the GWAS-removed data (±100 kb flank removal), using nearest-gene annotation. Top: Biological Process; Bottom: Cellular Component. As with Suppl. fig. 7, only PCs with at least one significant enrichment term are shown.

**Supplementary figure 21.**
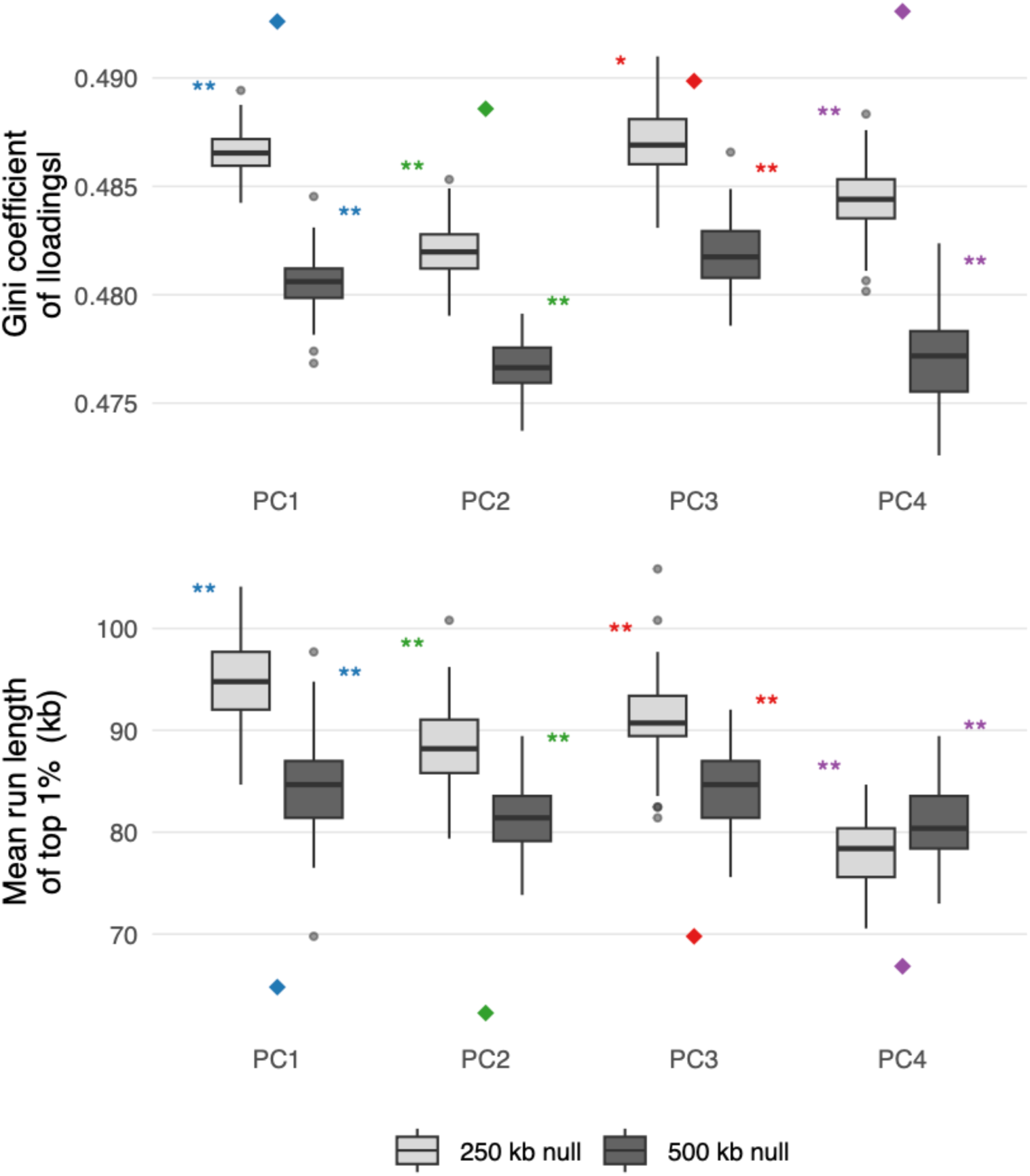
Within-block permutation null for loading concentration in the GWAS-removed PCA, PCs 1 – 4. This analysis is analogous to that in Figure S14 but permutations (100 total) and observed metrics are computed on the retained-window sub-matrix only. Significance values are all below 0.01.

**Supplementary figure 22.**
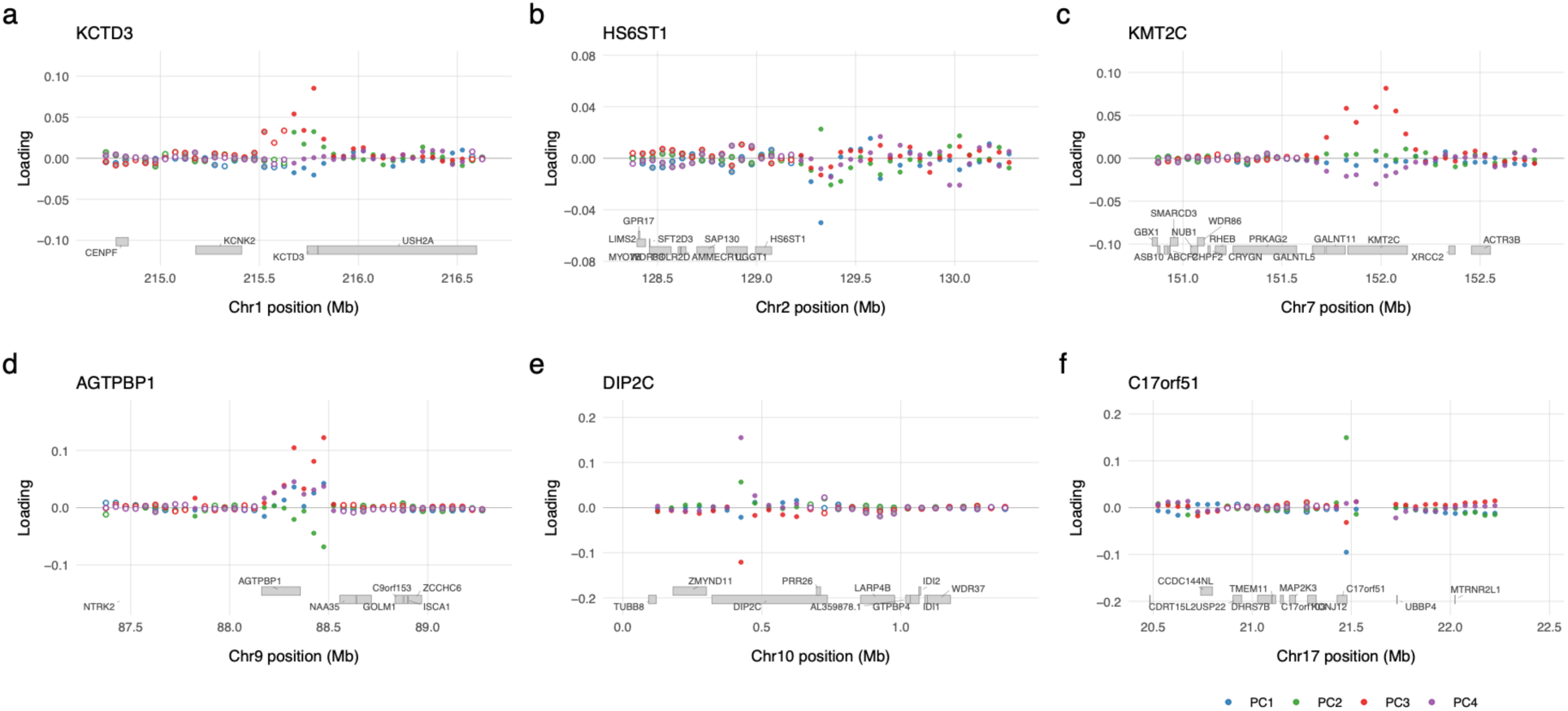
Zoomed regional plots for six regions with high PC loadings in the GWAS-removed data, analogous to Suppl. fig. 15. Each panel shows the loadings on PC1 – PC4, with the annotated genes for that region. Note that the direction of the loading is relatively arbitrary, and is a function only of the specific PC rotation selected. Empty circles indicate windows that were removed due to GWAS significance or presence in 100 kb flanking regions.

**Supplementary figure 23.**
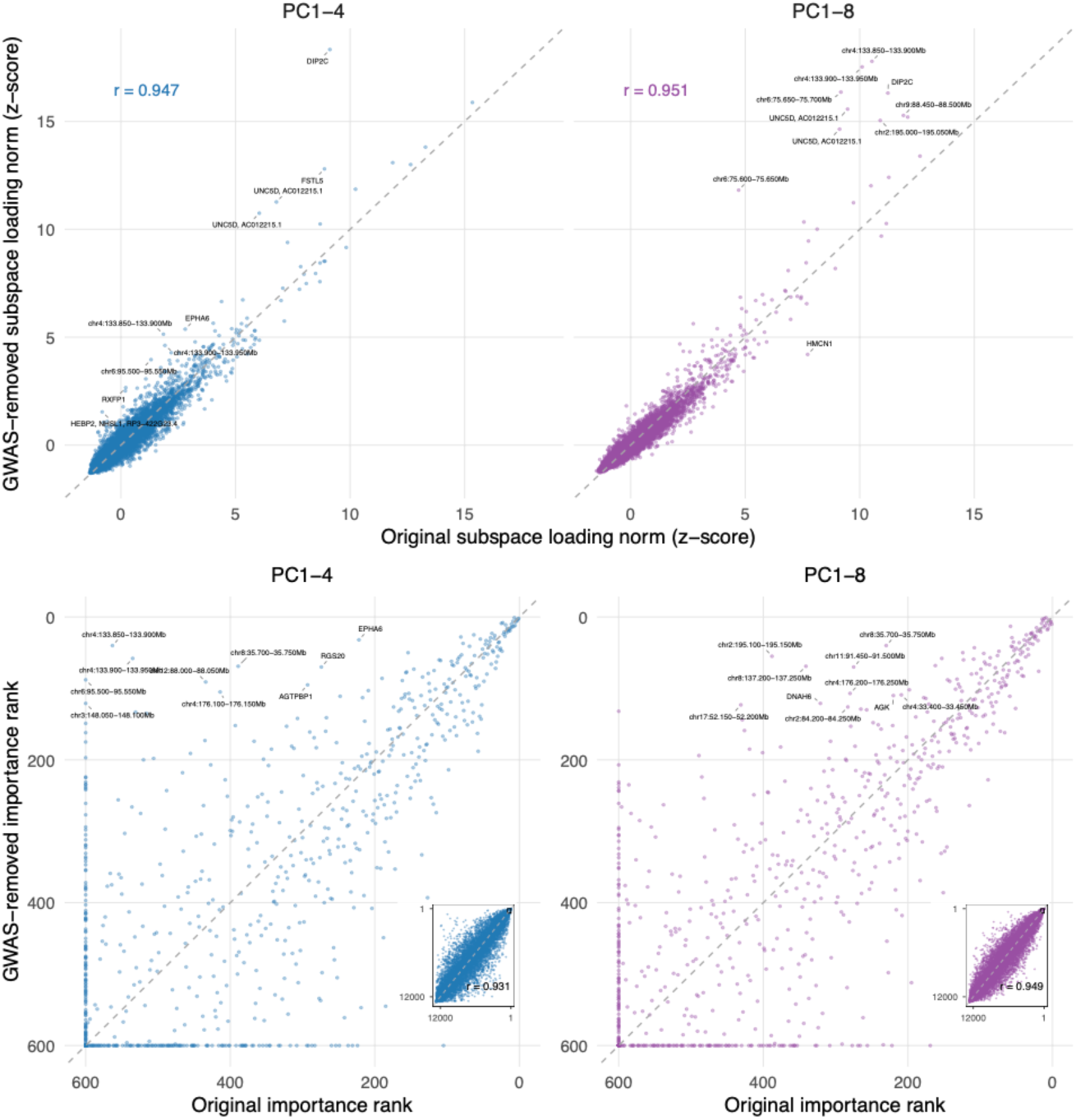
Comparison of per-window importance before and after removal of GWAS-significant loci (±100 kb flank-removal). Top row: Scatter of the sum of the absolute value of the loading weights for the PC1 – 4 (blue) and PC1 – 8 (purple). PC loading sums are z-score normalised, as loadings are generally heavier after GWAS-removal due to fewer available features. The dashed line indicates the x = y isocline. The 10 windows with the greatest change in PC loading sum are labelled with the overlapping genic annotation, or in case of no genic overlap, the chromosomal location. Pearson’s r is annotated at the top left. The loadings are heavily right skewed, as there are a relatively small number of loci with strong contributions. Bottom row: Rank scatter of the sum of the absolute value of the loading weights for the PC1 – 4 for the top 600 windows after GWAS removal. Note that axes are reversed (rank 1 is at the top right) to simplify interpretation. The dashed line indicates the x = y isocline. As in the top two panels, the 10 windows with the greatest change in ranking are labelled with the overlapping genic region, or in case of no genic overlap, the chromosomal region. Windows with ranks less than 600 either before or after are shown as having rank = 600. Insets: the full rank distribution, with Pearson’s r indicated in the bottom right of the inset. These values show no right skew as they are ranks and not z-scores.

**Supplementary figure 24.**
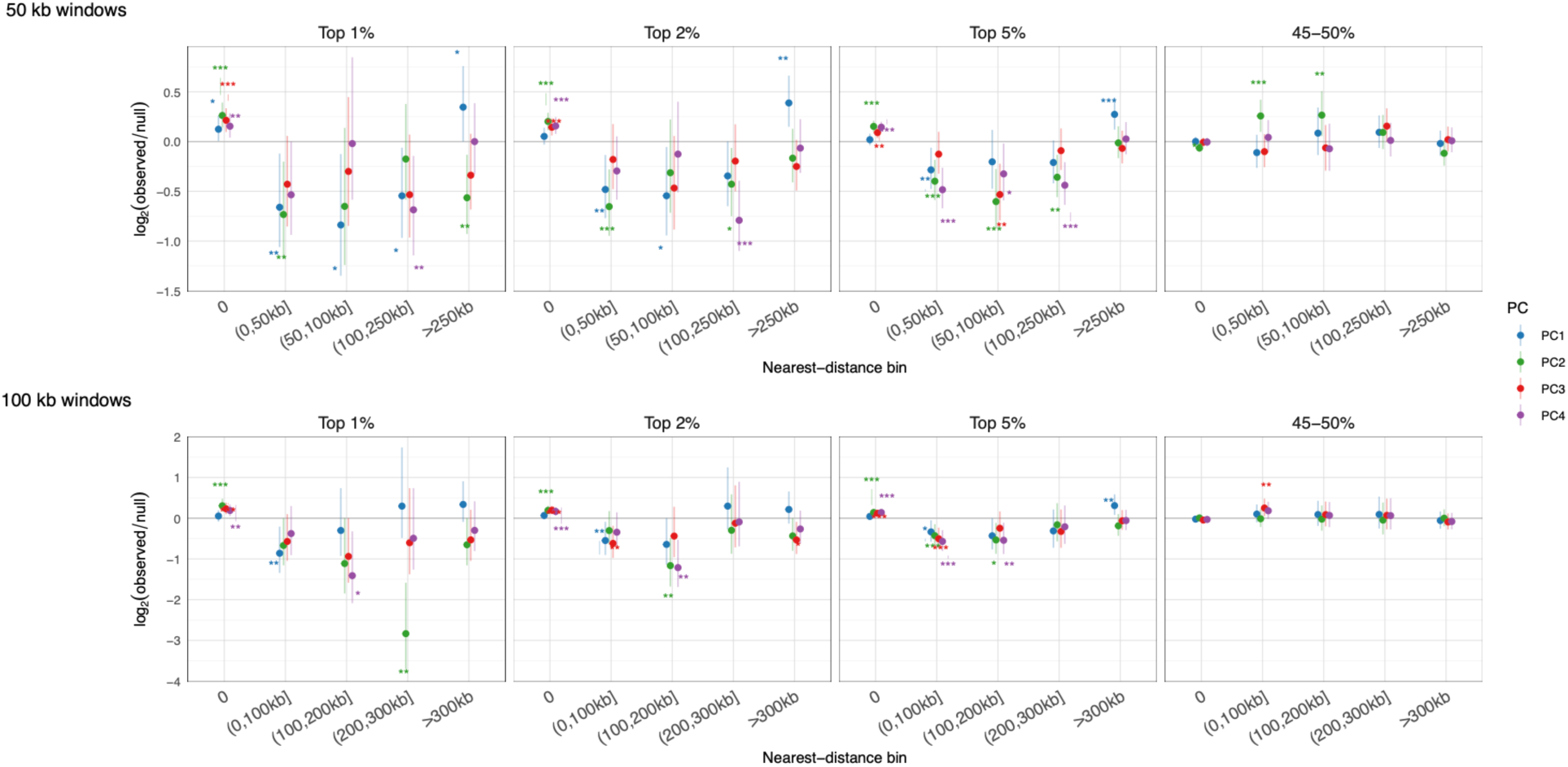
Nearest-gene distance bin enrichment (log₂ observed/null) for the top 1%, 2%, and 5%, most heavily loaded loci in the SVD for the full dataset, for both 50 kb and 100 kb window sizes. As with Suppl. Fig. 16 (which shows proximity to GWAS loci), the 45-50% bands are shown as a background permutation confirmation. Asterisks indicate significance: * p < 0.05; ** p < 0.01; *** p < 0.001.

**Supplementary figure 25.**
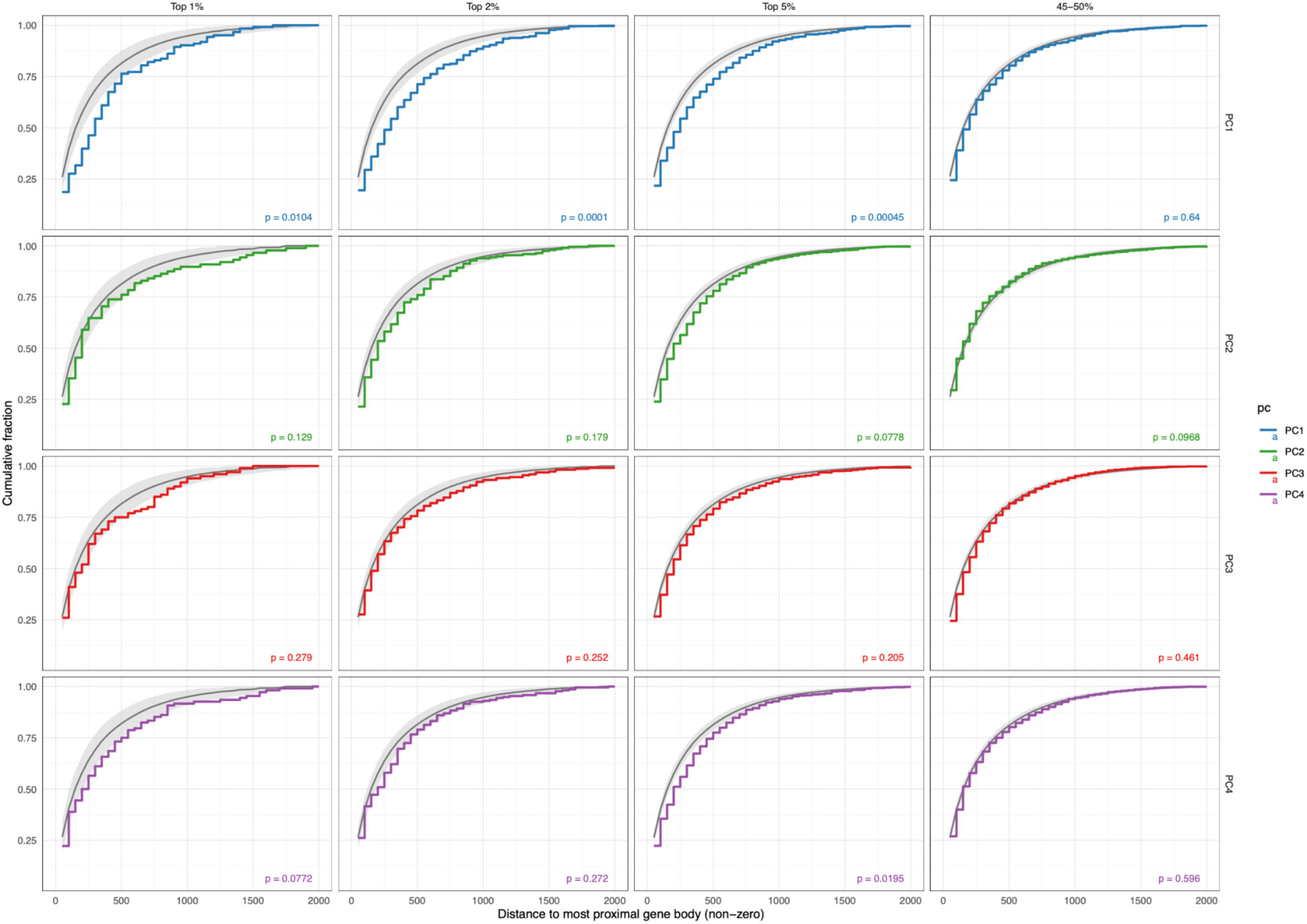
Cumulative distribution of nearest-gene distances, considering non-genic overlapping windows only, for top 1%, 2%, and 5% most strongly loaded 50 kb windows for PCs 1 – 4, with adjacent windows collapsed to a single window. Again, the 45–50% band is indicated as a background permutation control. Ribbons indicate the 2.5–97.5th percentile interval for 10,000 chromosome-matched null replicates; coloured steps are observed CDFs. PC1 shows consistent strong enrichment of loci distal to genic regions, with PC2 and PC4 showing less consistent gene-distal enrichment. p-values are the results of a chromosome-matched permutation test (10,000 replicates).

**Supplementary figure 26.**
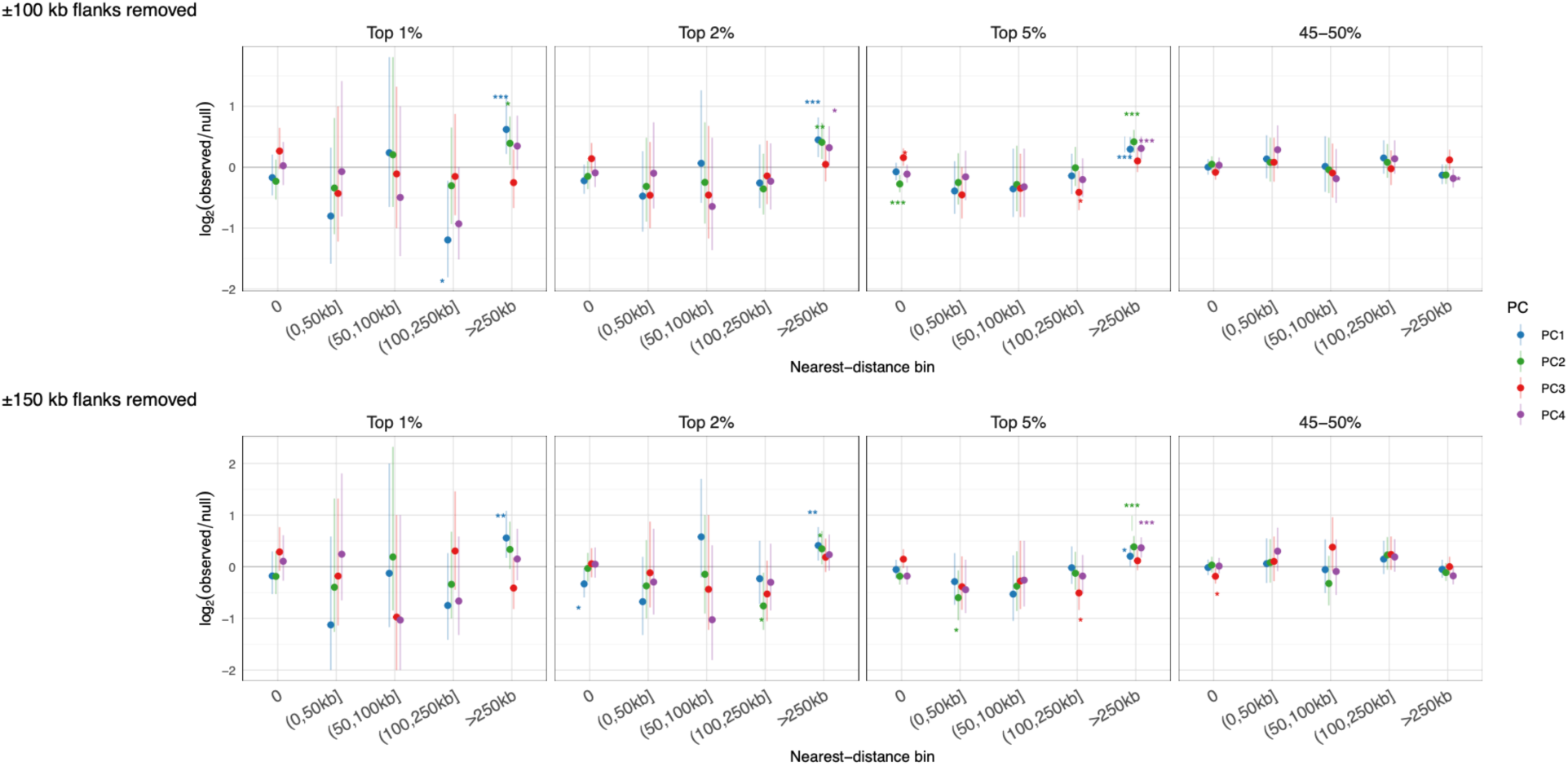
Nearest-gene distance bin enrichment (log₂ observed/null) for the top 1%, 2%, and 5%, most heavily loaded loci in GWAS-locus removed dataset, for both 100 kb and 150 kb flank removal. The 45-50% bands are shown as a background permutation control. In contrast with the results shown in Suppl. fig. 16, there is no consistent enrichment for gene-proximal windows, with a single exception: PC2 when considering the 5% most strongly loaded windows. However, there is consistent enrichment for gene-distal windows for PC1 (>250 kb), with weaker enrichment patterns for PCs 2 and 4. Asterisks indicate significance: * p < 0.05; ** p < 0.01; *** p < 0.001. This is analogous to Suppl. fig. 16 but for the GWAS filtered data.

**Supplementary figure 27.**
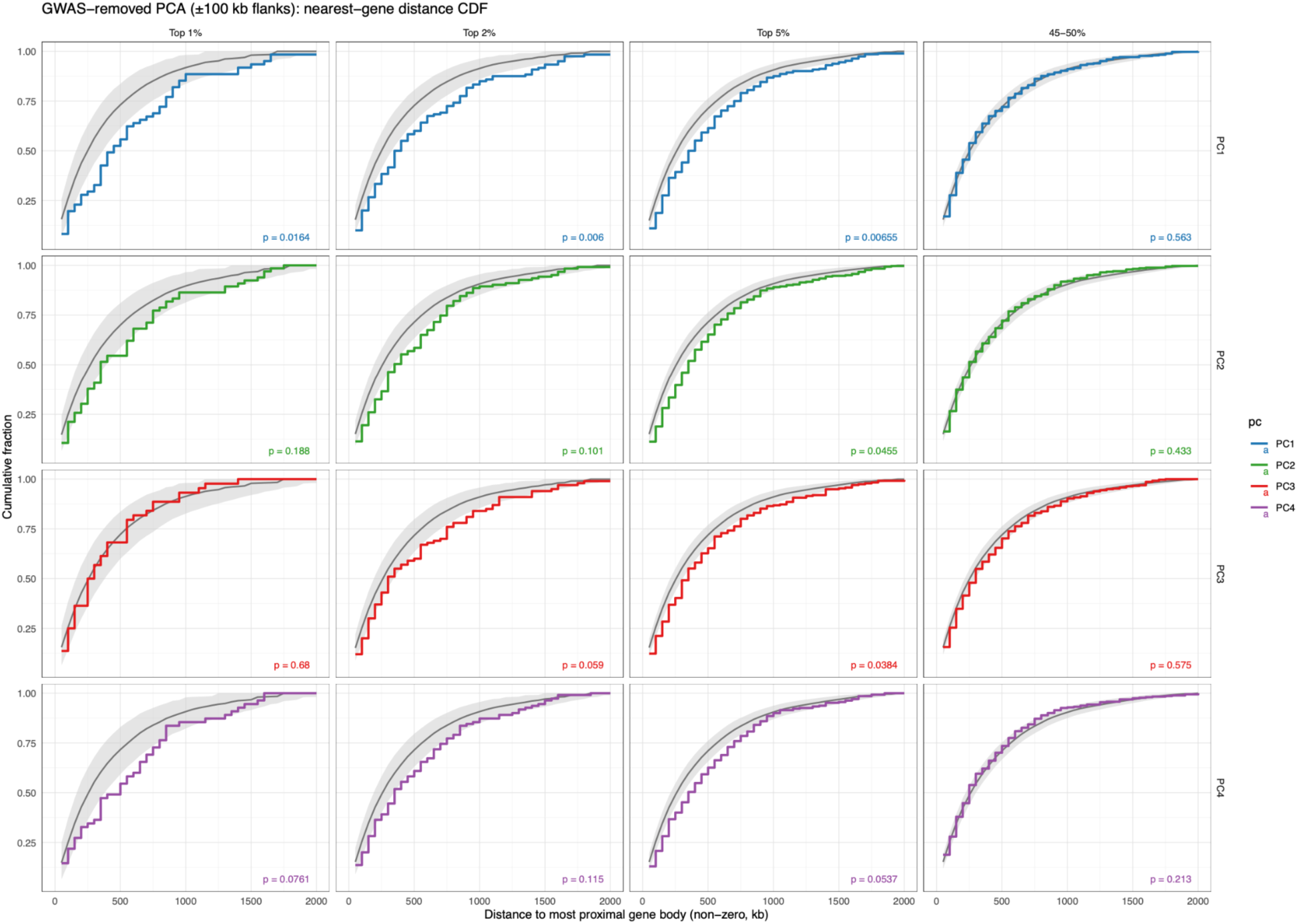
Cumulative distribution of nearest-gene distances considering non-genic overlapping windows only, for top 1%, 2%, and 5% most strongly loaded 50 kb windows for the GWAS-removed data, for PCs 1 - 4. Again, the 45–50% band is indicated as a background permutation control. Ribbons indicate the 2.5–97.5th percentile interval for 10,000 chromosome-matched null replicates; coloured steps are observed CDFs. p-values are derived from a chromosome-matched permutation test (10,000 replicates).

